# Drivers of genetic diversity across the marine tree of life

**DOI:** 10.1101/2025.06.03.657718

**Authors:** Rachel H. Toczydlowski, Reid S. Brennan, Eric D. Crandall, Joanna L. Kelley, James M. Pringle, Cynthia Riginos, John P. Wares, Gideon S. Bradburd

## Abstract

Why do some species have more genetic diversity than others? This question is one of the greatest remaining mysteries in evolutionary biology, yet we know little about what predicts genetic diversity among species. Efforts to quantify diversity and understand its ecological drivers have been hampered by a lack of genomic data processed in a consistent way and analyzed using statistical approaches that can move beyond the population level to generate species-wide estimates of genetic diversity. We address this critical gap by generating standardized estimates of genomic-level genetic diversity for 91 species sampled over 1,500 localities globally. Because marine species have evolved a striking diversity of biological traits that have long been hypothesized to affect species-level genetic diversity, we focus on marine taxa. For each species, we aggregate biotic traits related to life history and abiotic features of the species’ range. We show that species-level genetic diversity increases with species range extent and planktonic dispersal, and reason that these traits increase species’ demographic stability, thereby maintaining genetic diversity. Our findings provide insights into how biotic and abiotic factors interact to shape genomic variation in the ocean and offer a predictive framework for understanding marine biodiversity in the face of global change.

## Introduction

Why does genetic diversity vary among species? This is a central question in population genetics (Buffalo 2021; Leffler et al. 2012; Lewin & Eyre-Walker 2026; Lewontin 1974; Romiguier et al. 2014), and may inform a species’ adaptive potential, extinction risk, or conservation priority (Kardos et al. 2021; Mathur et al. 2023; Soulé 1992; Stange et al. 2021; Webster et al. 2017). Little is known about the factors that predict the observed variation in diversity across species (Ellegren & Galtier 2016). Indeed, for most species, we have no estimate of genetic diversity at all (Paz-Vinas et al. 2025; Toczydlowski et al. 2021).

Species-level genetic diversity, which we define as the total level of genetic polymorphism within a species, is ultimately the outcome of the relative rates of alleles entering and leaving the species gene pool and is thus controlled by population genetic and ecological processes that affect those rates, such as selection, mutation, and population demography (Ellegren & Galtier 2016). Although selection, including linked and background selection, can affect patterns of genetic diversity across the genome (Begun & Aquadro 1992; Elyashiv et al. 2016), evidence from both theoretical and empirical studies suggests that selection is unlikely to be the major determinant of variation in genetic diversity among species (Buffalo 2021; Coop 2016). And, while data on mutation rates are rare for most non-model organisms (Bergeron et al. 2023; Popovic et al. 2024), previous work suggests that variation in mutation rate alone is also unlikely to explain patterns of genetic diversity across species (Lewin & Eyre-Walker 2026; Romiguier et al. 2014, although see, e.g., Lynch 2010; Lynch et al. 2023).

Population genetic theory offers several hypotheses about the mechanisms by which different factors should increase or decrease the amount of genetic diversity present within a species. First, the genetic diversity of a population is expected to increase with the census number of reproductive individuals (Kimura & Crow 1963). Second, bottlenecks, whether local (i.e., confined to a local deme) or range-wide, are expected to decrease a species’ genetic diversity (Alcala & Vuilleumier 2014; Nei et al. 1975; Pannell & Charlesworth 1999). Third, dispersal is predicted to play conflicting roles; on the one hand, dispersal homogenizes genetic variation that has become locally differentiated due to genetic drift or selection (Slatkin 1985), and therefore can reduce the total diversity found across all demes of a structured metapopulation (Kimura & Weiss 1964; Maruyama 1972; Wilkins 2004). On the other hand, dispersal can help metapopulations avoid, or mitigate the severity of, bottlenecks (Stacey & Taper 1992), potentially increasing species-level genetic diversity in more dispersive species. Dispersal, in the form of increased migration, can also help save genetic diversity from being lost when the variance in fitness among demes is high, for example in the case of source/sink dynamics, where sinks rely on strong migration from sources to persist (Wang & Caballero 1999; Whitlock 2004; Whitlock & Barton 1997). Given these predictions from theory, it is clear that traits that impact a species’ population size, demography, and structure (or connectivity), have the potential to drive patterns of genetic diversity across species, although in directions that may not always be obvious (Lawrence & Fraser 2020).

There has been considerable research on patterns of genetic diversity across species and the factors that predict it (see Table S1), but much of this work suffers from limitations that affect the accuracy and generalizability of the results. For example, Leffler et al. (2012) reported that marine and freshwater chordates appear to exhibit higher genetic diversity than their terrestrial counterparts, and that species with larger geographic ranges tend to have higher genetic diversity than those with smaller ranges within the Drosophila genus (where data were rich enough to test this relationship); however, these results were generated from exploratory analyses without phylogenetic correction. Similarly, Romiguier et al. (2014), in a study of metazoan transcriptomes, found traits related to parental investment to be predictive of species-level genetic diversity, with r-selected species (i.e., high-fecundity, short-lived, small body-size) having higher genetic diversity than K-selected species, but only corrected for phylogeny at the inter-family level, and ignored the potential role of the geography of their sampling in driving observed patterns. Lewin & Eyre-Walker (2026) explicitly incorporate phylogeny into their analyses of factors predicting species-level effective population size across eukaryotes, and find that *Ne* decreases with generation time, propagule size, and adult body mass, and increases with their estimate of census size (product of population density and range size). However, they take no account of population structure or heterogeneity in the geography of sampling across taxa. Perhaps more importantly, this study is also a meta-analysis (rather than a synthesis) and therefore relies on published estimates of genetic diversity. These estimates are generated at the end of complex, multi-step, often idiosyncratic, bioinformatic pipelines which introduce noise (Hemstrom et al. 2024). Of compounding concern, if researchers share and recycle bioinformatic approaches more regularly within research communities coalesced around similar taxa, the bioinformatically introduced noise in estimates of genetic diversity may have a phylogenetic signal. In summary, understanding why levels of genetic diversity vary within and across species has received considerable attention (Clark & Pinsky 2024; De Kort et al. 2021) since the first molecular measurements of genetic variation were made (Hubby & Lewontin 1966; Lewontin & Hubby 1966), but despite these theoretical predictions and hypothesized relationships, performing a rigorous empirical test of these ideas has proven challenging.

Several factors have hindered empirical progress in understanding drivers of genetic diversity across species. First, quantifying genetic diversity remains difficult, particularly in species for which field sampling is laborious or costly. Estimates made from limited genetic information, such as mitochondrial DNA or microsatellites, might give noisy (Patterson et al. 2006) or biased (Birky et al. 1983; Rand 2001) estimates of diversity, but generating genomic datasets remains expensive. Second, even with genomic sequence data in hand, there are myriad bioinformatic pipelines and choices used to transform raw sequence data into assembled loci and called genotypes. Each bioinformatic decision point can have large impacts on downstream analyses (Hemstrom et al. 2024). To make meaningful comparisons of genetic diversity, it is therefore vital that the methods used for analyzing sequencing data and subsequently estimating genetic diversity are standardized across species, a bioinformatics challenge that has rarely been achieved at scale for genomic data. The findings generated by meta-analyses that fail to account for heterogeneity in bioinformatic data processing might be particularly sensitive to artifacts. Finally, in comparative population genetic studies, estimates of genetic diversity are implicitly treated as species-level traits, which requires correcting for phylogenetic nonindependence (Felsenstein 1985). Implementing such a phylogenetic correction necessitates an accurate, time-calibrated phylogeny that includes all species under consideration. Our understanding of phylogenetic relationships across the tree of life continues to grow, but many species still lack a well-supported branch tip, constraining the scope of comparative population genetic analyses that can be conducted.

Heterogeneity in sampling effort across species, and specifically in the spatial pattern and extent of sampling, poses another hurdle to a comparative analysis of species-level genetic diversity. Because many (if not most) species exhibit a pattern of isolation by distance (Meirmans 2012; Wright 1943) – as well as possible hierarchical or cryptic spatial structure – the geographic extent of sampling may often be a strong predictor of the estimated genetic diversity in a species. That is, a sample that covers a larger portion of a species range will encompass more intra-specific divergence and therefore diversity. Even if sampling effort were standardized, variation in density, dispersal, and recent demography across species’ ranges could result in biased estimates of a species’ genetic diversity (Exposito-Alonso et al. 2022). These factors impede our ability to accurately quantify genetic diversity within species and test hypotheses about its predictors across species.

In the context of global environmental change and the sixth mass extinction event currently occurring (Ceballos et al. 2015), as well as from a basic science perspective, the ability to determine the effect of different factors on genetic diversity is critical (Blanchet et al. 2017; Crandall et al. 2023; Paz-Vinas et al. 2018; Toczydlowski et al. 2021). Here, we begin to address this long-standing challenge in evolutionary biology by building a predictive framework using a novel, unified, spatially and phylogenetically explicit approach to test hypotheses about drivers of genetic diversity. We focus on marine biodiversity because oceans contain a taxonomically and functionally diverse biotic assemblage (Chaudhary & Costello 2023). How this marine biodiversity is generated and maintained in a dynamic, fluid environment, without obvious restrictions to gene flow, continues to be a fundamental problem in evolutionary biology (Hutchinson 1961; Knope et al. 2020; Palumbi 1994).

In addition to the unique complexities of the fluid ocean environment, marine biota exhibits tremendous variation in both range size and life history traits that govern connectivity across those ranges (Grosberg & Cunningham 2001). For example, marine biota encompass a great diversity of dispersal abilities (Chust et al. 2025; Pringle & Wares 2007; Sotka et al. 2004), and dispersal, particularly of drifting propagules, can be greatly affected by ocean currents (Becker et al. 2007; Riginos & Liggins 2013). Such interactions between biological traits and oceanic processes, both physical and chemical, generate spatial patterns of genomic diversity, which in turn might affect speciation and larger geographic patterns of species diversity (ÁlvarezNoriega et al. 2020; Chust et al. 2025; Ewers-Saucedo et al. 2016; Wares et al. 2001). Marine biota are also recognized as exhibiting considerable variation in population sizes (Buffalo 2021; McCauley et al. 2015; Steele et al. 2019). The diversity of traits present within marine organisms makes them well-suited for testing long-standing hypotheses about how biological traits and abiotic environments shape species-wide genetic diversity. There is also a critical need to understand and generalize such diversity patterns because oceanic biodiversity is being rapidly affected by climate change (Boyce et al. 2022; Pinsky et al. 2019; Poloczanska et al. 2013, 2016; Sunday et al. 2012) and other anthropogenic threats, including overfishing (Pinsky & Palumbi 2014), pollution (Rochman et al. 2015), disease (Harvell & Lamb 2020), and habitat change. Our approach addresses these gaps in understanding and gives insight into the drivers of genetic diversity across marine species while also establishing a template for future studies to test hypotheses about how and why genetic diversity varies across diverse taxa.

## Results

We generated the first species-wide, standardized estimates of genomic diversity for marine species worldwide and across the eukaryotic tree of life by leveraging a data synthesis approach and a novel spatially-explicit population genomic model. We first identified and downloaded all georeferenced, publicly available reduced-representation and whole-genome datasets of marine species present in the International Nucleotide Sequence Database Collaboration as of October 2020. We included as broad a phylogenetic diversity of species as we could in order to maximize the generality of our findings, but we excluded datasets containing laboratory-reared, freshwater, or captive populations, as well as those with individuals sampled outside their native range. The final metadataset of 91 sampled species covered a substantial proportion of the marine phylogenetic diversity of the eukaryotic tree of life (Figure 1), including representatives from vascular plants, invertebrate animals — including cnidarians, crustaceans, and molluscs — and vertebrate animals — including cartilaginous fishes, mammals, birds, reptiles, and ray-finned fishes. These datasets also met our constraint that they contain a minimum of 10 individuals (mean: 102, range: 10*−*366) sampled from at least 3 distinct geographic locations (mean: 17, range: 3*−*117). Starting from raw sequence reads, we filtered, cleaned, and assembled each of these 91 genomic datasets using multiple parameter combinations in a standardized bioinformatic framework, then picked the best set using an objective optimality criterion (see *Methods*), dropping low-coverage individuals. This resulted in a final genomic metadataset of 91 species (Figure 1A); each species was represented by a mean of 102 individuals (range: 8*−*366), 3.6 million raw sequence reads per individual (range: 0.4*−*30.1 million), 3.7 million aligned nucleotide base pairs across assembled genomic datasets (range: 5,800*−*22.3 million), and 37,017 loci (range: 86*−*187,685). To our knowledge, this represents the largest collection of interoperable co-analyzed population genomic datasets to date, terrestrial, freshwater, or marine.

**Figure 1.**
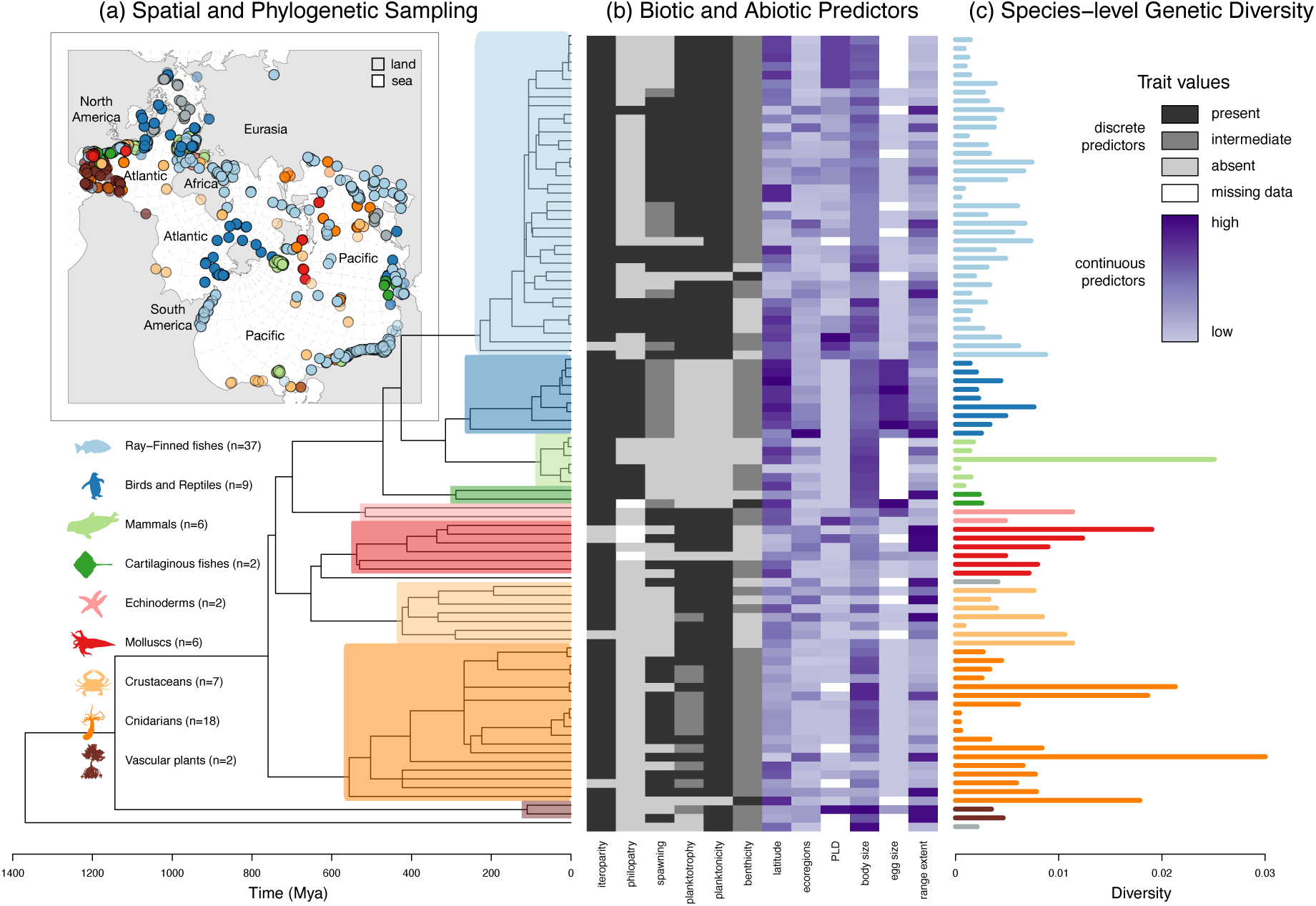
Species-level genetic diversity varies by almost two orders of magnitude in the world’s oceans. (A) Time-calibrated phylogeny of all species (*N* = 91) and map of all samples (*N* = 9,325) included in this study. The map uses the Spilhaus projection which, by emphasizing the oceans and their connectivity, rather than land masses, represents the world as a marine organism might see it. (B) Set of discrete (gray-scale) and continuous (blue-scale) biotic and abiotic predictors collated for each species. Colors represent the trait state or value for each species. (C) Species-level genetic diversity estimated using a spatially explicit model and genomic sequence data.

We then employed the novel parametric modeling approach introduced by Hancock et al. (2024) and Hancock & Bradburd (2025) to generate estimates of species-level genetic diversity that are robust to heterogeneity in sampling area and intensity. Briefly, we fit an asymptotic curve to the observed Isolation by Distance (IBD) pattern of pairwise homozygosity (Malécot 1969; Wright 1943) in each species. We calculated geographic distance here as the distance traveling overwater between the locations of sampled individuals; pairwise homozygosity is the probability that a pair of alleles sampled at random from two individuals within the species are the same. The asymptote of this IBD curve — a parameter estimated as part of the model-fitting procedure — captures the extrapolated genetic diversity contained within the entire species range, which we refer to as *π*. By including multiple individuals per species and estimating diversity in an explicitly geographic statistical framework, this approach represents an advance on previous estimates made from a small number of “representative individuals” that are treated as experimental replicates. Our estimates of *π* in the world’s oceans varied greatly across species, spanning almost two orders of magnitude from 3.371 *×* 10*^−^*^4^ to 3.001 *×* 10*^−^*^2^, with a mean of 5.284 *×* 10*^−^*^3^ (phylogenetically corrected mean: 5.311 *×* 10*^−^*^3^) (Figures 1C and S1).

To test hypotheses about which biotic and abiotic traits predict genetic diversity across marine species, we assembled a time-calibrated phylogeny and aggregated biotic trait data as well as abiotic factors for all 91 species in our dataset (Figure 1B). Genetic diversity had significant phylogenetic structure (Pagel’s *λ*=0.832, *p* = 5 *×* 10*^−^*^4^; Blomberg’s *K*=0.106, *p* = 0.019); the correlation between phylogenetic relatedness and similarity in genetic diversity was significant over the last 490 My, and the strength of the correlation increased monotonically as it approached the present (Figure S2). Genetic diversity is also affected by species-level characteristics and mechanisms that emerge as the result of the interaction of biotic and abiotic processes (Jablonski 2008), such as reproductive census population size, the frequency, magnitude, duration, and geography of bottlenecks (i.e., where in a species’ range bottlenecks occur and how large of an area they affect), and effective dispersal ability. These characteristics and mechanisms are difficult to directly observe, so instead we focused on biotic and abiotic traits that are hypothesized to affect them (Table S1), and for which sufficient published data were available (>70% of the species in our analyses; Figure S3). The suite of biotic traits is comprised of body size, egg size, spawning, pelagic larval duration, philopatry, iteroparity, planktotrophy (larval feeding behavior), benthicity (degree of association with seafloor), and planktonicity (qualitative association with drifting development as a component of dispersal). Abiotic predictors consisted of mean species’ range latitude (Figuerola-Ferrando et al. 2023; Pegan et al. 2025; Wares et al. 2021), the number of ecoregions within the species’ range (ecological complexity and its potential to act as a barrier to gene flow), and overall range extent. We note that, because traits evolve in the shared context of the lineages in which we measure them, they may frequently occur as part of a “syndrome,” i.e., a set of traits or trait values that tend to co-occur either due to constraints or selection. These macroevolutionary syndromes may make it difficult to isolate the effect of a particular predictor on species-level genetic diversity. We expand on this point in the Discussion, and all traits are further described in the *Supplemental Methods* and Appendix S1.

Range extent, number of ecoregions, and planktonicity had significant positive effects on species-level genetic diversity (Figures 2 and 3); species with planktonic life stages, as well as species with larger range extents or ranges that included more ecological regions, all had higher expected *π*. We give the estimated effect sizes of the biological parameters in Table S2, and the estimated effect sizes of the “nuisance” parameters in Table S3. Range extent, defined as the greatest overwater distance between any pair of individuals of that species in the Global Biodiversity Information Facility (GBIF), with filtering to ensure incidental range-edge observations did not inflate this value, had a mean effect of 0.52 (95% CI: 0.016–0.96). Number of ecoregions, defined as the number of ecoregion polygons from Spalding et al. (2007) that contained filtered GBIF observation points for that species, had a mean effect of 0.006 (95% CI: 1.12 *×* 10*^−^*^5^ – 0.012). We use ecoregions to quantify the number of discontinuities in biodiversity and abiotic features within a species’ range that may limit gene flow (and thereby increase species-level genetic diversity). Planktonicity, defined as whether the species was planktonic at any point in their life cycle, had a mean effect of 0.580 (95% CI: 0.032–1.19). Planktotrophy and free-spawning had a strongly positive estimated effect on genetic diversity, and the effects of body size, iteroparity, and philopatry were strongly negative. However, these effects’ credible intervals overlapped zero and were not inferred to be significant (see *Methods*), so those results should be interpreted with caution. Range extent and number of ecoregions were highly correlated (Kendall’s *τ* = 0.78; Figure S4), and subsequent analyses (see *Methods*) suggest that the inferred effect of ecoregions on genetic diversity is at least partially driven by the range extent. Planktonicity and range extent were very weakly correlated (Kendall’s *τ* =-0.06; Figure S4), indicating that the significance levels of their effects are largely independent. The Bayesian *R*^2^ values (Gelman et al. 2019) for the models that included range extent and planktonicity as biological predictors (Figure 3) were 0.56 (95%CI: 0.32-0.76) and 0.56 (95%CI: 0.33-0.76) respectively.

**Figure 2.**
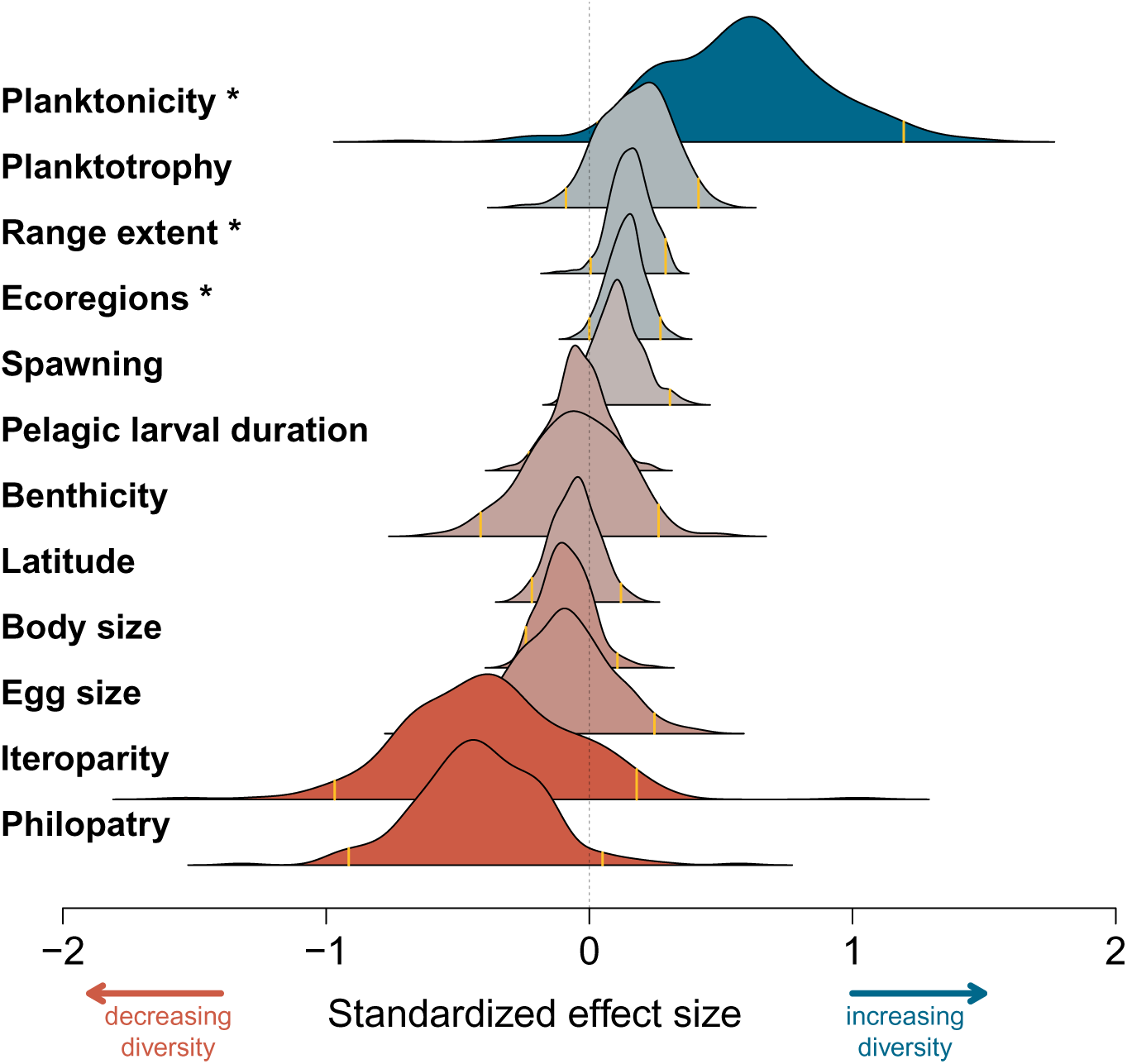
The estimated effects of biotic and abiotic predictors on species’ genetic diversity vary in their magnitude and direction. Estimated effect sizes (presented via their marginal posterior distributions) are standardized by the standard deviation of their predictor and are ordered from top to bottom by mean effect on species-level genetic diversity. Significant predictors – those for which the 95% equal-tailed credible interval of the marginal distribution of effect sizes did not overlap zero (gray, vertical dashed line) - are denoted with an asterisk.

**Figure 3.**
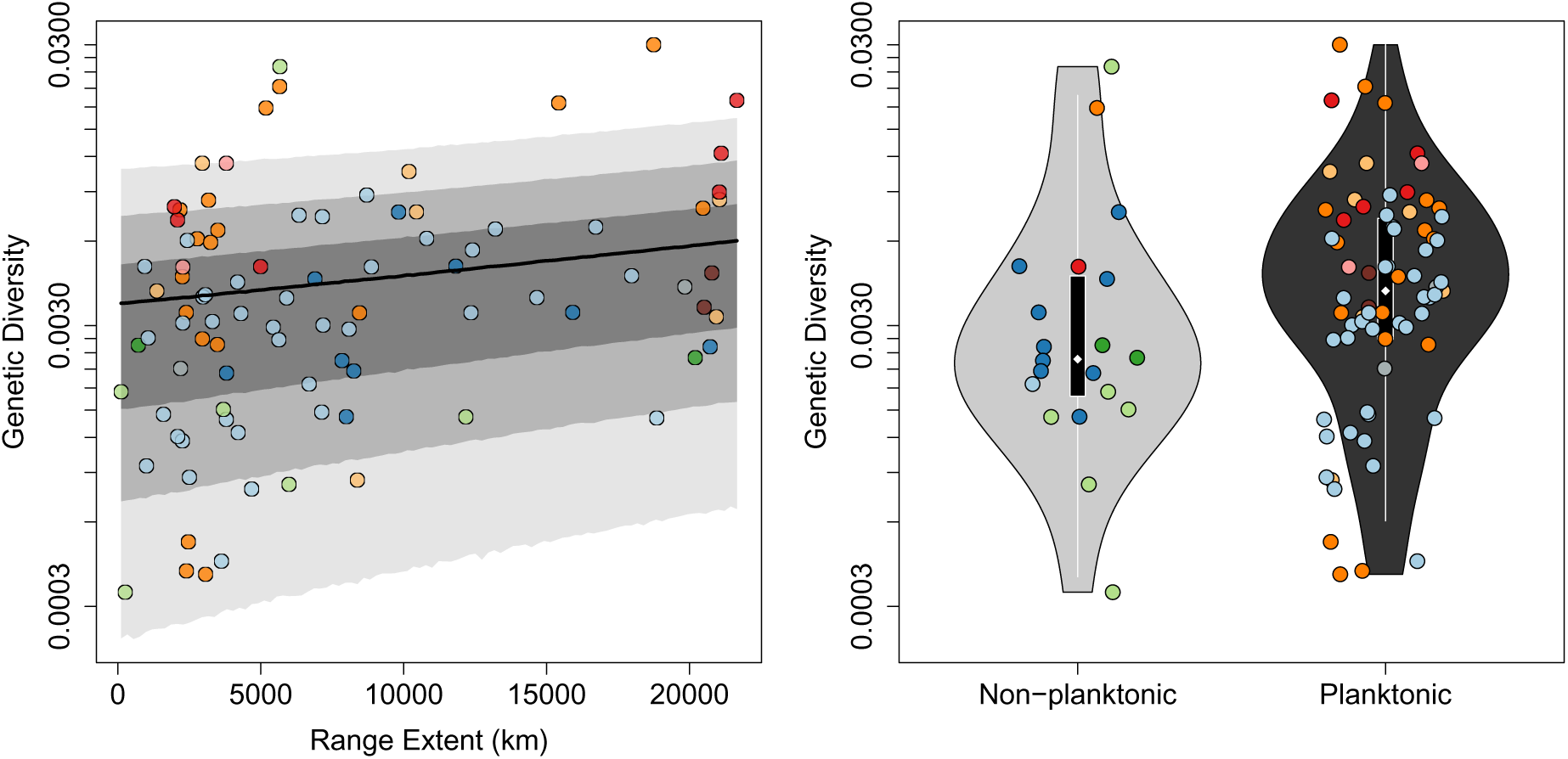
Species-level genetic diversity is most significantly predicted by species range extent and the presence of a planktonic life stage. (A) Species with larger range extents have higher genetic diversity. Shaded polygons represent 50%, 80%, and 95% model prediction credible intervals. Mean Bayesian *R*^2^ = 0.56 (95%CI: 0.32-0.76). (B) Species that have at least one planktonic life stage have higher genetic diversity than species that are never planktonic. Mean Bayesian *R*^2^=0.56 (95%CI: 0.33-0.76). Each point represents a unique species (*N* = 91) and is colored by taxonomic clade (as in Figure 1).

## Discussion

Our analyses reveal substantial variation in the amount of genetic diversity harbored within marine species across the tree of life. Estimates of species-level genetic diversity varied over almost two orders of magnitude. Such variation offers immense opportunity to test hypotheses about what factors predict genetic diversity across species. We found strong phylogenetic signal in genetic diversity, suggesting that diversity may be reasonably predicted in species of known phylogenetic position but lacking population genomic sampling resources. We also found strong evidence that range extent and planktonicity were positively correlated with genetic diversity (Figure 3), in line with previous findings about the roles of range size and dispersal (Leffler et al. 2012; Lewin & Eyre-Walker 2026; Romiguier et al. 2014). We found that the majority of dispersal-related traits yielded positive effects on genetic diversity, despite the role that dispersal can play in decreasing metapopulation diversity by homogenizing local variation. We hypothesize that these relationships reflect the effects of dispersal-related traits on both species’ capacity to avoid or recover from bottlenecks as well as in the maintenance of diversity that might otherwise be lost due to among-deme variance in population dynamics (e.g., sources/sinks). Below, we walk through our results (including predictor effect sizes that were not significant but nonetheless had considerable estimated posterior mass far from zero) and explore their possible interpretations in light of population genetic theory and previous empirical research.

Based on population genetic theory, the genetic diversity of a population is expected to increase with its census size of reproductive individuals (Kimura & Crow 1963). In lieu of estimates of species census size, which would be unreliable for the majority of taxa included in our study, we included two factors likely to be correlated with census population size: body size and range extent. Body size is expected to be negatively correlated with census size (Brown 1995) and we find that body size has a negative effect on genetic diversity (although we note that the effect of body size on genetic diversity may be mediated by correlations between body size and other biotic traits, such as dispersal (Chust et al. 2025)). Conversely, range extent is expected to be positively correlated with census population size (James & Eyre-Walker 2020; Rosenfield 2002), and, consistent with predictions from theory, we find a positive effect of range extent on diversity (Figure 2). Although neither body size nor range extent is a perfect predictor of population census size, their mean estimated effect sizes are nonetheless consistent with the predicted effects of census size on diversity.

Range extent is expected to be correlated with census size, and thereby with species-wide genetic diversity, but it can also influence a species’ genetic diversity via its impact on spatial population genetic processes (Maruyama 1972; Wilkins 2004). This is because lineages separated by a larger distance must wait longer before coalescing, and the geographic distance between a pair of lineages can be larger in a larger area. Thus, larger ranges increase the expected time to most recent common ancestor of a sample and thereby also increase the amount of standing genetic diversity. Ecological complexity within a range is also expected to increase the overall amount of genetic diversity, as ecological heterogeneity and discontinuities in abiotic features and biotic communities may act as direct barriers to dispersal or decrease effective migration due to selection against locally maladaptive alleles and linked variation (Sobel et al. 2010; Wang & Bradburd 2014). However, because the total amount of ecological complexity contained within a range is inextricably linked to the range size (e.g., Figure S4), it is difficult to tease apart the relative contributions of range extent and ecological complexity to total genetic diversity. In addition, species with larger range sizes may be more likely to conceal cryptic species (Cahill et al. 2023), the existence of which would artificially inflate their estimated diversity.

The majority of dispersal-related traits were estimated to have positive effects on species-level genetic diversity. Under a model with low variance in the demographic contributions of different demes to the metapopulation, the opposite pattern is predicted: dispersal homogenizes local genetic variation (that arises due to genetic drift, selection, and mutation) across the demes of the metapopulation. Therefore, dispersal is expected to reduce the total amount of genetic differentiation – and thereby genetic diversity – found collectively across the demes in a spatially structured metapopulation (Kimura & Weiss 1964). However, when variance in deme-level demographic contributions is high – for example, when demes experience local bottlenecks or persistent source/sink dynamics – increased dispersal is expected to reduce the rate of genetic drift in the metapopulation, increasing species-level genetic diversity (Wang & Caballero 1999; Whitlock 2004; Whitlock & Barton 1997).

In lieu of direct estimates of the effective parent-offspring dispersal distances (Bradburd & Ralph 2019) for the species in this study, we used biological traits with demonstrated relationships to effective dispersal: planktonicity, planktotrophy, spawning (internal vs. nesting vs. free-spawning), pelagic larval duration (PLD), benthicity, and philopatry (Table S1). Planktonicity, planktotrophy, and more dispersive spawning modes were positively correlated with species-level genetic diversity, while philopatry was negatively correlated. Note that plank-tonicity and planktotrophy were strongly correlated with each other (Kendall’s *τ* = 0.76), and both were negatively correlated with philopatry (*τ* =-0.60 and-0.38 respectively; Figure S4), so it is possible that some of the signal in the estimated effects of these predictors are due to those correlations. We also acknowledge that planktonicity and planktotrophy are not perfect predictors of dispersal ability (e.g., even direct developers can disperse long distances (Spencer et al. 2025)). However, in general, planktonic dispersal and longer planktonic durations lead to longer dispersal distances (Pringle 2023; Shanks 2009) and there is abundant evidence that the mean potential for dispersal is substantively greater for planktotrophs than non-planktotrophs (Chust et al. 2016; Schiebelhut et al. 2022b; Wares et al. 2001).

For both of the remaining dispersal-related traits, PLD and benthicity, the majority of the posterior mass of the estimated effect size was centered on zero, suggesting these traits may be poor predictors of species-level genetic diversity. We expect PLD may be poorly quantified for many species given the variation we observed in the literature both in estimation techniques and reported values within species. As for benthicity, our results suggest that whether or not adults eventually attach to a substrate may have less bearing on species-level genetic diversity as compared to the dispersive capacity of their gametes and offspring (note only three species in our dataset are benthic their entire lives, providing little power to test this syndrome). In sum, the fact that the majority of the dispersal traits were positively correlated with diversity (and none had strong negative effects) is consistent with the hypothesis that the ability of dispersal to mitigate genetic drift outweighs its homogenizing effect.

The associations of dispersal-related traits with a species’ ability to recover quickly after a local bottleneck, or to maintain sink populations via the net export of migrants from sources, might explain why their estimated effects were the opposite of what is predicted via dispersal’s direct effect on diversity. Population bottlenecks will reduce the genetic diversity of a species, to an extent that is determined by their longevity and severity (Nei et al. 1975), so species traits that reduce the prevalence or severity of bottlenecks, or that increase the rate of recovery after a bottleneck, are predicted to increase species-level genetic diversity. Traits associated with higher dispersal (e.g., planktonicity, planktotrophy, dispersive spawning modes) might lead to faster recovery from local bottlenecks, as extirpated demes may be recolonized more quickly if there is greater connectivity among demes (Schiebelhut et al. 2022a,b; Villarino et al. 2018). And, conversely, philopatry, which decreases the connectivity between demes, might lead to slower recovery from local bottlenecks and decreased genetic diversity (Kawecki & Holt 2002; Waples 2025).

Other life history traits are also predicted to affect the capacity of a species to recover from a bottleneck, and thereby genetic diversity. Species with higher fecundity are also expected to recover more quickly from bottlenecks and thus to harbor more genetic diversity. In their comparative population genomic analysis, Romiguier et al. (2014) and Lewin & Eyre-Walker (2026) found strong support for this hypothesis, with more fecund species harboring more genetic diversity, as well as those species with smaller propagules. In line with this result, we found egg size (which is often inversely correlated with fecundity (Kasimatis & Riginos 2016; Marshall & Keough 2007; Romiguier et al. 2014)) to have a negative inferred effect on diversity. However, unlike previous work in this area (Lewin & Eyre-Walker 2026; Romiguier et al. 2014), our fecundity proxy (egg size) was not statistically significantly correlated with genetic diversity. Interestingly, iteroparity, which could buffer local demes from drops in genetic diversity due to bottlenecks, especially bottlenecks resulting from a generation of failed reproduction (Clark et al. 2024; Ellner & Hairston Jr 1994), had a strong estimated negative effect on predicted diversity. This could be due to non-causal associations with other traits likely to decrease genetic diversity (e.g., census population size, age at sexual maturity). Iteroparous species are also more likely to have longer generation times and lifespans, both of which have been shown to be predictive of decreased diversity (Lewin & Eyre-Walker 2026; Waples et al. 2013).

Lastly, while dispersal and fecundity are expected to affect species’ responses to bottlenecks, geographic position may influence how frequently species experience bottlenecks. Populations closer to the equator are hypothesized to experience greater abiotic stability (Rohde 1992; Willig et al. 2003) and may therefore be subjected to fewer extreme climatic events and associated bottlenecks. In addition, due to their geological history, lower latitude regions have had more similar abiotic environments for longer (Brodie & Mannion 2023). Both patterns have been proposed as explanations for the latitudinal gradient in species diversity (Fine 2015; Lawrence & Fraser 2020, but see Chaudhary et al. 2016). These hypotheses also suggest predictions for *intra*-specific diversity: namely that genetic diversity should be higher in lower latitudes (Clark & Pinsky 2024). Consistent with this prediction, we find that species with lower mean range latitudes have higher genetic diversity.

There are a number of important caveats to our results. First, the lack of information on variation in mutation rate across taxa precluded employing mutation rate as a predictor in our models. However, assuming mutation rate is phylogenetically correlated (Drummond et al. 2012; Ramos-Almodovar et al. 2025), variation in genetic diversity across species due to variation in mutation rate will have been captured by the phylogenetic covariance structure of our regression analyses. Second, the presence of unidentified cryptic species could inflate our estimates of genetic diversity as both allelic diversity and heterozygosity will be overestimated in these cases (Pante et al. 2015; Riginos et al. 2024). We treated each species or subspecies as identified in the original published papers as a distinct taxon in our analyses, including when estimating species-level genetic diversity; we believe that the authors of the original studies are best suited to identify and delineate species in this case. Nevertheless, species delimitation was beyond the scope of our analyses and it is possible cryptic species are present within datasets we treated as a single species. Cryptic species are found in both vertebrates (Zhang & Wiens 2026) and invertebrates (Li & Wiens 2023), though the frequency varies among taxa (PérezPonce de León & Poulin 2016). However, this signal again is likely phylogenetically correlated and may therefore be partly accounted for in our analyses. For example, in the marine environment, corals are recognized as having a high prevalence of cryptic species (Riginos et al. 2024). Third, it is possible that our findings may not generalize to terrestrial species. There are many features of marine life that are not shared with terrestrial species, including the advective medium and the prevalence of gamete or propagule dispersal via that medium (Steele et al. 2019). An exciting area of future research would be to expand our analytical approach to terrestrial species and test whether range size or dispersal-related traits are also predictive of species-level genetic diversity in this realm. Fourth, we focused on deep evolutionary time, yet the lability of many life history traits can be quite high even within families or genera (Hart et al. 1997). Therefore, we hope to see complementary comparative population genomic analyses that focus on more recent phylogenetic scales. Only a minute subset of marine diversity has been studied intensely enough to support such analyses (Dietrich et al. 2020; Paz-Vinas et al. 2025), but this will change as large-scale datasets become increasingly available. Finally, though we have controlled for phylogenetic dependence on the associations studied, still greater insights could be gained from removing as much of the phylogenetic effect as possible (Zakas et al. 2018).

## Conclusions

No single factor explained the majority of variation in genetic diversity across taxa (unlike, e.g., Romiguier et al. (2014)). Instead, we found two factors that were significantly predictive of genetic diversity in the world’s oceans; species with larger range extents had higher genetic diversity, as did species with at least one planktonic life-stage. The effects of other predictors, though not significant, suggest that species traits that facilitate greater resilience to, or quicker recovery from, bottlenecks have higher diversity. These results should be digested with the caveat that we cannot rule out the possibility that some portion of the estimated effect sizes for these predictors may be driven by their correlations with other predictors in our dataset or with other traits not included in our study.

Our results also suggest tantalizing connections between the microevolutionary and macroevolutionary patterns of variation. One exciting future avenue of research would be to explore how, or whether, factors that predict high genetic diversity at the species level are also predictive of high diversification rates at the clade level, although we note that predictions here are by no means clear (Januario et al. 2025; Riginos et al. 2014).

Taken together, our work sheds light on the mystery of heterogeneity in genetic variation across species and suggests intriguing future avenues of comparative genomic research. We hope to see the marine genomic and trait database that underpins this study grow and facilitate such lines of inquiry in the research community moving forward. Additionally, the novel analytical approaches that we used establish a template for robustly testing hypotheses about what drives genomic-level diversity across diverse taxa. Predicting spatial patterns of diversity based on life history traits and spatial distribution represents a significant step forward in both fundamental research and applied conservation.

## Materials and methods

### Locating genomic datasets

We located and downloaded all georeferenced, publicly available, reduced-representation and whole-genome sequence datasets of marine species present in the International Nucleotide Sequence Database Collaboration (INSDC) as of October 2020. We included an additional 48 datasets for which we retrieved sampling coordinates external to INSDC (Crandall et al. 2023). We retained datasets that had *≥*10 georeferenced individuals and *≥*3 sampled locations (Figure S5). We excluded samples from non-native ranges, freshwater/inland locales, captive or lab populations, quantitative genetic studies, ancient time, pooled genetic sequences, known species hybrids, individuals only identified to genus, and salmonids (due to philopatry to freshwater habitats). We confirmed that none of the final species we analyzed exhibit philopatry to freshwater (as this behavior is not unique to salmonids).

### Bioinformatic processing of genomic datasets

Starting from the raw sequence reads, we filtered, cleaned, and assembled each genomic dataset *de novo* using a standardized, custom pipeline and Stacks2 (Rochette et al. 2019). Post-Stacks, we dropped low coverage individuals (mean number of genotyped and/or co-genotyped base pairs *<*30% of median for dataset) and low coverage loci (scored in *<*50% of remaining individuals). We then used the r80 method to pick the optimal *de novo* assembly for each dataset of nine different standardized Stacks2 parameter combinations tested (Paris et al. 2017); we used the optimal assembly for each dataset (unique species) to estimate species-level genetic diversity.

### Estimating species-level genetic diversity

We estimated species-level genetic diversity for each of the 91 species in this study using a Bayesian parameterization of the Wright-Malécot isolation by distance model (IBD) (Hancock & Bradburd 2025; Hancock et al. 2024). This approach allowed us to generate comparable estimates of genetic diversity across datasets that differed in sampling extent (number of individuals, absolute geographic area, and proportion of species’ range). We used pairwise homozygosity between sequenced individuals and pairwise sea-wise distances (the distance in kilometers between sampling locations traveling via water and avoiding land) as model inputs. We used this sea-wise geographical distance, which is agnostic to ocean currents, as a marine version of Isolation by Distance. Implementing a distance metric that includes the effects of ocean currents (e.g., Jahnke & Jonsson 2022; White et al. 2010) would necessitate incorporating information about PLD, planktonicity, and benthicity (Treml et al. 2015), which would have generated collinearities that confounded our ability to test the effects of these biotic traits on species-level genetic diversity. It is also unclear how ocean currents would be relevant to dispersal and connectivity in nektonic species.

We ran five independent chains of the Wright-Malécot IBD model per dataset and reported results from the chain with the highest mean log posterior probability (5,000 iterations/chain, 2500 iteration burn-in, chains thinned to every 10th iteration). In all downstream analyses, we used an estimate of *π* (post burn-in mean of the marginal distribution from the best chain) as our species-level estimate of genetic diversity. This parameter is best understood as long-term diversity; in a spatially structured population, it describes the expectation of the maximum genetic divergence between any pair of samples in a species, regardless of where they are sampled in the species range. This quantity is related to the long-term effective population size, and is determined by the species mutation rate, dispersal rate, and the geometry of its range (Wilkins 2004).

### Collecting biotic, abiotic, and phylogenetic predictors

We collected a suite of biotic traits, abiotic traits, and phylogenetic divergence times for all species in the study to test the extent to which they predict species-level genetic diversity. We selected biotic traits hypothesized to influence levels of genetic diversity via their associations with population size, bottlenecks (i.e., degree of demographic stability), and/or dispersal (Table S1). These include: iteroparity (whether individuals can mate multiple times before death, or once); philopatry (habitual return to birthplace to mate); spawning mode (i.e., internal vs. nesting vs. free-spawning, which relate to the degree of parent-progeny dispersal); benthicity (degree to which the adult species is associated with the ocean floor); planktonicity (whether the species has a planktonic stage); planktotrophy (larval feeding behavior, which is related to early-life dispersal and rvs K-selection); pelagic larval duration (PLD time spent drifting in the plankton); egg size; and body size. See Appendix S1 for further descriptions of how these biotic traits were defined.

We tested abiotic predictors related to the size and ecological complexity of each species’ range, which may influence species-level genetic diversity via associations with population size and levels of genetic divergence. Specifically, we quantified: range extent (maximum linear extent of range); mean range latitude (absolute value of mean latitude of range); and the number of unique ecoregions present within each species’ range (Marine Ecoregions of the World (Spalding et al. 2007)). We used species occurrence records from the Global Biodiversity Information Facility (GBIF) (GBIF: The Global Biodiversity Information Facility 2020) as input for estimating these three abiotic predictors.

Finally, we used the January 2022 beta release of TimeTree 5 (Kumar et al. 2022) to obtain estimates of the pairwise divergence times between all 91 species in the study. For the 24 species that were not present in TimeTree, we assigned them randomly to a tip within the clade representing the lowest/shallowest taxonomic level possible or formed a polytomy.

### Modeling effects of predictors on genetic diversity

We tested for an effect of phylogenetic relatedness on similarity in *π* using Blomberg’s *K* (Blomberg et al. 2003) and Pagel’s *λ* (Pagel 1999) as implemented in the *phytools* R package (Revell 2024). We tested for a correlation across evolutionary time between phylogenetic relatedness and similarity in *π* using a phylocorrelogram (Keck et al. 2016) and used a permutation approach to assess significance. We tested hypotheses about the effects of biotic and abiotic predictors on *π* using a custom Bayesian phylogenetic multiple regression model implemented in Rstan (Stan Development Team 2024). Our response variable, *π*, is bounded between 0 and 1, so we model it with a Beta distribution, using the mean/precision implementation of the Beta (Ferrari & Cribari-Neto 2004). The mean of the Beta distribution is connected to the phylogenetic multiple linear regression via a logit link function. See the subsection of the *Supplemental Methods* titled “Modeling effects of predictors on genetic diversity” for full model details. To reduce model complexity, avoid issues with multicollinearity, and increase the interpretability of estimated coefficients, we estimated the effect of each biotic and abiotic factor on *π* one at a time, in each case incorporating a phylogenetic variance-covariance structure (Garland & Ives 2000) as well as four technical “nuisance” predictors: (1) number of individuals in the species’ dataset; (2) the mean raw read count; (3) the mean read length; and (4) the mean locus depth.

By including these nuisance predictors in each analysis and jointly estimating their effects with those of the biotic and abiotic factors in which we are interested, we ensured that technical artifacts are not having a significant impact on the results of this study. We calculated the Bayesian *R*^2^ (Gelman et al. 2019) of each model (including the four nuisance parameters and each of the biological parameters in turn).

## Acknowledgments

This research is the product of a working group formed at the National Science Foundation Research Coordination Network for Evolution in Changing Seas Synthesis Workshop (NSF-OCE 1764316, awarded to Katie E. Lotterhos). We thank the authors of Toczydlowski et al. (2021) and Crandall et al. (2023) for locating geospatial coordinates that were absent in INSDC for 48 of the genomic datasets included in this study. We are grateful to Bruce Martin for writing code to assist with phylogenetic analyses and the Spilhaus map projection. Ricardo Lemos also provided code for implementing the Spilhaus projection. Pat Bills, Yaniv Brandvain, Doc Edge, Steve Goldstein, Morgan Kelly, Katie Lotterhos, Bruce Martin, Misha Matz, Marjorie Weber, Will Wetzel, Mike Whitlock, Ben Winger, Max Witynski, and members of the Bradburd lab all provided helpful feedback and scientific discussion. Computational work was performed using computational resources and services provided by the Institute for Cyber-Enabled Research at Michigan State University.

## Statement of authorship

All authors contributed to conceptualization and writing. R.H.T. and R.S.B. designed the bioinformatics workflow and processed the genomic datasets. E.D.C., C.R., and J.P.W. amassed and curated biotic trait data. J.M.P. and R.H.T. calculated abiotic predictors. G.S.B. conceived of population genetic analyses. R.H.T. and G.S.B provided supervision and leadership. We hold the time we had to work with and learn from E. D. Crandall fondly and with gratitude; more about his life and career can be read at https://rcn-ecs.github.io/.

## Data accessibility

All data have been previously published (See Appendices S3, S5, and S6 in Supplemental Information for citations). All code is archived in a public git repository (permanent version upon time of publication of this manuscript: doi:XXXXX; dynamic version: https://github.com/gbradburd/divdiv).

