## Supplemental Materials for "Drivers of genetic diversity across the marine tree of life"

**This PDF file includes:**

- Supplementary Methods
- Tables S1 to S3
- Figures S1 to S9
- Appendices S1 to S6

### Supplementary Methods

#### Locating genomic datasets

We searched the International Nucleotide Sequence Database Collaboration (INSDC) to locate publicly available genomic-level sequence data for marine organisms. We accessed the INSDC from the United States National Center for Biotechnology Information (NCBI) data center using R (R Core Team 2023) and the *rentrez* R package (v1.2.3, (Winter 2017)). We used the entrez_search function to locate entries in INSDC and entrez_summary and/or entrez_fetch to download the associated metadata. We first identified all non-human, non-viral, non-metagenomic, and non-bacterial datasets (BioProjects) that were focused on DNA sequencing and had publicly available sequence data. Next, we downloaded all metadata for the individuals in each of these datasets from the Sequence Read Archive (SRA) and BioSample databases. We retained sequences with library strategies: WGS, WCS, WGA, RAD-Seq, or OTHER. We then queried the remaining organism names (assigned in the BioSample database) in the World Register of Marine Species (WoRMS 2021 Editorial Board 2021) to identify marine organisms. We used the *worms* R package (v0.2.2, (Holstein 2018)) with no fuzzy name matching to execute this query.

We then filtered the remaining potentially marine genomic datasets to those that included at least 10 unique individuals (BioSamples) and 3 unique collection locations (based on the geospatial coordinates provided in the BioSamples database). Finally, we manually dropped samples (or datasets) from non-native ranges, captive or lab populations, quantitative genetic studies, ancient time, and pooled genetic sequences. We also dropped known species hybrids, salmonids (due to philopatry to freshwater habitats), and freshwater/inland samples. In cases where there were multiple genomic datasets for a species, we retained the dataset with the broadest geographic sampling extent (as our goal was to estimate species-level genetic diversity). We manually recovered latitude/longitude from associated scientific publications for 48 datasets that met all of our filtering criteria but lacked spatial coordinates in INSDC (Crandall et al. 2023); we dropped those where latitude/longitude were unrecoverable. We performed this search strategy on October 28, 2019 and again on October 19, 2020. See Appendix S2 for the exact search strategy and Appendix S3 for the citations of all 91 genomic datasets used in this study.

#### Bioinformatic processing of genomic datasets

We started with the raw reads published in the SRA for each relevant dataset, then cleaned, filtered, and assembled them into loci using a standardized bioinformatic pipeline. Applying the same bioinformatic steps to all datasets allowed us to make meaningful comparisons across datasets. Furthermore, many studies associated with the publication of these datasets only published diversity metrics (and/or derived files) for polymorphic loci. As a result, we could not calculate absolute measures of diversity without re-assembling the reads to obtain the number of both polymorphic and nonpolymorphic base pairs sequenced (Korunes & Samuk 2021). The following steps were all performed on the Michigan State Institute for Cyber-Enabled Research High Performance Computing Cluster (https://icer.msu.edu/).

We downloaded raw reads from the SRA using the fasterq-dump function in the SRA Toolkit v2.10.7 (Kodama et al. 2012). We treated all paired-end datasets as single-end datasets, retaining all forward reads (regardless of whether they had a reverse pair) and discarding all reverse reads

(SRA Tools fasterq-dump –split-files option). Next, we used the Stacks2 process_radtags module v2.54 (Rochette et al. 2019) to remove reads with uncalled bases and/or where the mean Phred score in a sliding window 15% of the read length dropped below 15. After removing low quality reads, we used cutadapt v2.1 (Martin 2011) to search for and remove adapter sequences (e.g. the Illumina common adapter) from the 3' end of reads. See Appendix S4 for the complete list of next-generation sequence adapters that we searched for. Datasets with adapters were easy to identify by the presence of adapter hits in >25% of the reads (compared to ∼5% of reads in clean datasets). About a fifth of the datasets (20 of 91) contained adapters. We confirmed that no adapter sequences remained in cleaned, trimmed reads by running a final adapter search. Lastly, we trimmed all reads to a standardized length within each dataset; *de novo* loci assembly is more robust when all raw reads are a uniform length within each dataset (Rochette et al. 2019). For datasets with variable length reads, we selected a uniform read length for each dataset by visually assessing the distribution of read lengths for the dataset and choosing a length that retained long reads in a majority of samples.

We assembled each dataset *de novo* using Stacks2 v2.54 (Rochette et al. 2019). For each dataset, we identified a set of optimal assembly parameters following guidelines from (Paris et al. 2017) and (Rochette et al. 2019). We used a custom parallel workflow to assemble each dataset with 9 different parameter combinations of *M* (ustacks) and *n* (cstacks): $\{M = 3, n = 2\}$, $\{M = 3, n = 3\}$, $\{M = 3, n = 4\}$, $\{M = 6, n = 5\}$, $\{M = 6, n = 6\}$, $\{M = 6, n = 7\}$, $\{M = 9, n = 8\}$, $\{M = 9, n = 9\}$, and $\{M = 9, n = 10\}$. These are the parameters that most strongly influence assembly behavior (Rochette et al. 2019). The *M* parameter controls the number of mismatches allowed between reads within individuals and *n* controls the number of mismatches allowed between loci among individuals. We used default values for all other parameters.

For each assembled dataset and parameter combination, we dropped individuals with a mean number of genotyped and/or co-genotyped base pairs <30% of the median for the dataset (including nonpolymorphic and polymorphic sites). These individuals with few final assembled loci also commonly had few raw reads (suggesting they were poorly sequenced, rather than poorly assembled). We then dropped any loci that were scored in less than 50% of the remaining individuals. We performed this filtering after the populations module using the populations.samples.fa and populations.snps.vcf Stacks output files and custom R scripts. Finally, we selected the assembly parameter combination (of the nine standardized combinations tested) that yielded the highest number of polymorphic loci scored in at least 80% of individuals for each dataset (r80 method, (Paris et al. 2017)). The r80 method has been demonstrated to identify the set of assembly parameters that yield the maximum number of conservatively plausible polymorphic loci for a genomic dataset.

#### Estimating species-level genetic diversity

Estimates of species-level genetic diversity can be heavily influenced by the number of individuals sampled, where in the species' range those samples are located, and what proportion of the species' range is sampled. Such biases result from spatial genetic structure (i.e., correlations between genetic ancestry and geographic location) within a species' range (Battey et al. 2020; Bradburd et al. 2018; Exposito-Alonso et al. 2022). To generate unbiased estimates of diversity from datasets with heterogeneous sampling, we used the Bayesian parameterization of the Wright-Malécot model of isolation by distance (IBD) (Malécot 1969; Wright 1943) implemented in Hancock & Bradburd (2025); Hancock et al. (2024) for each of the 91 species in our study. Briefly, this method implements the theoretical model of isolation by distance introduced by Wright (1943) and Malécot (1969) to describe the decay of genetic relatedness between individuals as an asymptotic function of the geographic distance separating them. The asymptote – a parameter estimated by the model-fitting procedure – captures the mean maximum genetic divergence between individuals of a species, regardless of their geographic separation (Hancock et al. 2024). In Hancock *et al.*, the estimate of the value of this asymptote is designated as collecting phase $\pi$, or $\pi_c$, in reference to the collecting phase of the spatial coalescent (Wilkins 2004); for simplicity, we refer to it as $\pi$ throughout this manuscript. Our estimate of species-level genetic diversity is informative about long-term inbreeding $N_e$ and relatively insensitive to recent demography (Hancock et al. 2024). Because this approach explicitly models the rate of isolation-by-distance, our estimates of species-level diversity are robust to both the number of samples and their spatial distribution (and comparable across studies with heterogeneous sampling schemes).

To calculate species-level genetic diversity, the Hancock et al. (2024) model implementation requires as inputs – a matrix of pairwise homozygosity (identity-by-state) and a corresponding matrix of pairwise geographic distances between genotyped individuals. We calculated pairwise homozygosity as 1-$D_{XY}$ using custom R scripts following Hancock et al. (2024); these scripts accounted for missing genotypes by defining the denominator for $D_{XY}$ as the total number of base pairs co-genotyped for each pair of individuals (within a dataset). We calculated pairwise geographic distance as the distance between each pair of individuals traveling via the sea (avoiding crossing land). We used the National Oceanic and Atmospheric Administration ETOPO1 global bathymetric data with a resolution of 4 arc-minutes (Amante & Eakins 2009) and the *trans.mat* and *lc.dist* marmap functions of the *marmap* R package v1.0.6 (Pante & Simon-Bouhet 2013) to calculate these sea-wise distances. For individuals that fell on land (e.g., near coastlines), we used the *dist2isobath* marmap function with isobath = 0 to relocate each "land point" to the closest point in water. We calculated each sea-wise distance using two resistance matrices (one with longitudes from -180 to 180, the other with longitudes from 0 to 360) and retained the shorter distance.

For each species dataset, we then ran five independent chains of the model in Hancock et al. (2024) for 5,000 iterations each. We discarded the first 2,500 iterations as burn-in and sampled the remaining chain every 10th iteration to reduce autocorrelation. We discarded chains that mixed poorly and identified the single best chain for each species as that with the highest mean log posterior probability. We used the mean of the marginal distribution of $\pi$ from the best chain for each dataset as our estimate of species-level genetic diversity in downstream analyses.

To quantify variation in diversity across populations within species, we also calculated $\pi$ for all individuals sampled at each unique geographic location included in the sampling dataset. We report the standard deviation of those location-specific estimates of $\pi$ in Figure S6, and we show the relationship between variance in diversity and our global estimates of diversity for each species in Figure S7. The comparison between variation in location-specific estimates of diversity and the species global estimate shows a positive correlation, which is an artifact of the mean-variance relationship.

#### Collecting biotic, abiotic, and phylogenetic predictors

**Biotic predictors** We focused on nine biotic traits related to reproduction and dispersal: philopatry, iteroparity, spawning, egg size, benthicity, planktonicity, planktotrophy, pelagic larval duration

[PLD], and body size. We selected these traits because they have demonstrated associations with intraspecific spatial genetic diversity and divergence or are expected to be correlated with genetic diversity via their associations with population size, bottlenecks, and/or dispersal (Table S1). We defined each discrete trait (i.e., all but PLD, egg size, and body size, which are continuous) as "present," "intermediate," or "absent" for each species (Figure 1).

Specifically, we defined iteroparity as "present" for species that can breed multiple times per lifetime, and "absent" for species that can only breed once. We defined philopatry as "present" for species where individuals return to their natal breeding area to reproduce and "absent" for species that do not perform such migrations. We defined three categories for spawning to capture variation in how far embryos disperse from parents. Embryos can be internally fertilized and remain attached to the parent ("absent"), laid in nests or other benthically attached structures that are often (but not always) dissociated from the parents ("intermediate"), or free-floating, typically the product of broadcast spawning ("present"). Note this is a distinct life history facet from the presence or absence of planktonic larvae (see below). We defined the trait benthicity to characterize the degree to which species have a relationship with the ocean floor (or other fixed location). Benthicity was "present" for species that maintain a strong connection to the bottom throughout their lifetime (e.g., hydrozoans and anemonefishes that lack planktonic larvae), "absent" for species that are never associated with the bottom (e.g., dolphins that give birth at sea to swimming young), and "intermediate" for species that have both benthic and pelagic life stages (e.g., fish with planktonic larvae and benthic adults). We used planktonicity to distinguish species that have at least one planktonic – floating or drifting – life stage ("present") from those that are never planktonic ("absent").

Planktotrophy is a predictor of the amount of time larvae spend drifting in the water column feeding, which is related to the level of provisioning provided by the egg as well as the capacity for external feeding. We coded planktotrophy as "absent" for species that lack a (planktonic) larval stage and instead develop directly from embryo into young that resemble miniature adults. We coded planktotrophy as "intermediate" for species that produce planktonic larvae that feed on yolk from well-provisioned eggs to facilitate metamorphosis into juveniles (i.e., lecithotrophic larvae); as these larvae are planktonic, they drift in the water column while developing, but develop relatively quickly given the well-provisioned eggs. Lastly, we coded planktotrophy as "present" for species that produce planktonic larvae that must feed on particulates in the water column to fuel metamorphosis into juveniles given they derive from minimally-provisioned eggs (i.e., planktotrophic species); development usually takes longer than planktonic larvae derived from well-provisioned eggs (Strathmann 1987). We defined pelagic larval duration (PLD) as the mean (or midpoint) number of days that marine larvae spend in the plankton. Finally, we defined egg size as the largest dimension of the egg at hatching (or progeny at birth) and body size as the largest dimension of an adult individual. Extended trait definitions are available in Appendix S1.

When possible, we used trait values reported in FishBase (R. Froese, D. Pauly, Editors 2023), SeaLifeBase (M. L. D. Palomares, D. Pauly. Editors 2023), and large compilations (e.g., Cowen & Sponaugle 1997, for reef fishes). For most values, however, we performed Boolean searches in Web of Science or Google Scholar (e.g., <species name> AND egg size). We were able to obtain trait values for $> 90\%$ of the species in our study for all traits except egg size (73% coverage across species; Figure S3). All species had values for $> 77\%$ of the nine biotic traits (Figure S8). Full citations for all biotic trait values are listed in Appendix S5.

**Abiotic predictors** We used three abiotic predictors to characterize the geographic extent and amount of ecological complexity encompassed by each species' range – range extent, mean range latitude, and the number of ecoregions encompassed within the species' range. We estimated these three quantities for each species using species occurrence records from the Global Biodiversity Information Facility (GBIF) (GBIF: The Global Biodiversity Information Facility 2020). We first downloaded all GBIF presence records for each species that were not fossils, not managed, and did not have identified geospatial issues using the occ_download() function of the *rgbif* R package (v3.5.0, (Chamberlain et al. 2017)) on September 8, 2021 (see Appendix S6 for exact search parameters and full citations for all downloaded GBIF data). We then visually examined maps of the occurrence records returned for each species and manually removed spatial outliers (that were unlikely to be correctly identified records) and observations within non-native ranges. We used a combination of literature, expert opinion, and published range maps to decide which additional GBIF occurrence records, if any, to manually drop for each species.

We defined the range extent as the longest axis of the species' range "as the fish swims," that is, traveling via the sea and avoiding crossing land. For each species, we calculated all pairwise "sea-wise distances" between the final, cleaned GBIF occurrence records using the same approach that we employed for the genotyped individuals (see *Estimating species-level genetic diversity*). We then used the value at the $97.5^{\text{th}}$ percentile of the resulting pairwise "sea-wise distance" distance matrix (between species occurrence records) as the final range extent; using the $97.5^{\text{th}}$ percentile minimized the influence of any one individual occurrence record, especially those near the range edges. For computational efficiency, we thinned occurrence datasets that had more than 15,000 points using a raster proportional to the spatial extent of the dataset before generating the pairwise "sea-wise distance" distance matrix.

We defined mean range latitude as the absolute value of the mean latitude of all the final, cleaned GBIF occurrence records for each species. We used original locations in this calculation, (rather than relocating "land points" to the nearest ocean), but mean latitudes were highly correlated between the two approaches (Pearson's $R$ = 0.98). We did not include the genetic sample locations in the range extent or distance from equator calculations because they were nested within the spatial extent of, and often duplicated in, the GBIF occurrence records.

Finally, we characterized the amount of ecological complexity contained within each species' range by calculating the total number of unique Marine Ecoregions of the World (MEOW) ecoregions that at least one GBIF occurrence record or genetic sample fell within (Spalding et al. 2007). We refer to this predictor as "number of ecoregions." MEOW ecoregions are biogeographic units, developed in large part to facilitate conservation priority setting and planning, whose boundaries reflect synthetic discontinuities in taxonomic assemblages, dominant habitats, ocean currents and temperature, and geomorphological features (Spalding et al. 2007). MEOW ecoregions classify coastal and shelf systems, where the majority of (known) marine species diversity resides (Spalding et al. 2007). Note, the number of ecoregions was highly correlated with the spatially coarser MEOW realms and provinces as well as the Costello et al. (2017) ecoregions (Pearson's $R$ all $>$ 0.87). Transitions between ecoregions may act as barriers to gene flow within a taxon, so even though a given species may not be found in every ecoregion contained within their range, the number of ecoregions may nonetheless be predictive of total population genetic diversity of the species.

**Phylogenetic relationships** We derived estimates of the pairwise divergence times between all species in the study using the January 2022 beta release of TimeTree 5 (Kumar et al. 2022). We first identified species that were present in TimeTree (67/91 species in study). When a focal species was not in TimeTree but other closely related species were (more closely related than any other species in our study, based on published taxonomy), we assigned the focal species randomly to a tip within the clade representing the lowest/shallowest taxonomic level possible (17 species assigned to a representative tip in their genus, 1 species assigned at level of family). This approach is sensible because any tip within the focal clade will give the same divergence time to species outside of the clade, however, it assumes that the oldest species defining the focal clade is present in TimeTree. Divergence times for focal species in clades where this assumption is not met will be upwardly biased. Six focal species were missing from TimeTree and shared a clade with other species in this study (meaning we could not assign them to a random tip within the clade). For one of these six species, we were able to identify the most closely related species present in TimeTree (using published literature) and use this tip. For the remaining five species absent from TimeTree, we added a tip to the tree that formed a polytomy with the lowest taxonomic level of classification present in TimeTree.

#### Phylogenetic comparative analyses

We evaluated the strength and significance of the phylogeny in predicting patterns of similarity in diversity using three complementary approaches. We first estimated the overall significance of phylogenetic structure in genetic diversity using Blomberg's $K$ (Blomberg et al. 2003) and Pagel's $\lambda$ (Pagel 1999), both implemented with the function *phylosig* in the R package *phytools* 2.0 (Revell 2024). We tested for a correlation between phylogenetic relatedness and similarity in $\pi$ using the R package *phylosignal* 1.3.1 (Keck et al. 2016). We determined significance of correlation through time using a permutation approach: we first randomly permuted tip diversity values and, for each permutation, we estimated the correlogram using the *phylosignal* package. We repeated this permutation approach 500 times and used the equal-tailed 95% confidence interval of these "null" simulations to determine the significance of the observed correlogram for diversity.

#### Modeling effects of predictors on genetic diversity

To test hypotheses about the effects of biotic and abiotic predictors on $\pi$, we used a custom Bayesian phylogenetic multiple regression model with Beta-distributed errors (Ferrari & Cribari-Neto 2004) implemented in Rstan v2.32.3 (Stan Development Team 2024). Specifically, we used the mean/precision parameterization of the Beta distribution shown below:

$$\pi \sim \text{Beta}(a, b) \quad (1)$$
$$a = \mu \times \phi \quad (2)$$
$$b = (1 - \mu) \times \phi \quad (3)$$
$$\mu_i = \frac{1}{1 + \exp(-\theta_i)} \quad (4)$$
$$\vec{\theta} \sim \mathcal{N}\left(\gamma + \sum_M \beta_m \vec{X}_m \ , \ \alpha\Psi\right) \quad (5)$$

where $a$ and $b$ are the shape parameters of the Beta distribution, and $\mu$ and $\phi$ function as the mean and precision. The mean of the Beta distribution, $\mu$, is given by the inverse-logit transformed parameter $\theta$, which is modeled as a draw from a multivariate normal distribution. The mean of that multivariate normal distribution is given by the expression $\gamma + \sum_M \beta_m \vec{X}_m$, where $\gamma$ is a global mean of $\theta$, $M$ is the number of predictor variables, $\vec{X}_m$ is the vector of predictor variable values for the $m$th predictor, and $\beta_m$ is the effect size of the $m$th predictor. The covariance of the multivariate normal distribution on $\theta$ is given by $\alpha\Psi$, where $\Psi$ is a phylogenetic correlation matrix (i.e. a phylogenetic variance-covariance matrix scaled so that all values fall between 0 and 1), and $\alpha$ is the estimated Brownian rate parameter (i.e. the macroevolutionary heritability) (Garland & Ives 2000). We generated the phylogenetic variance-covariance matrix from our ultrametric phylogeny using the R package *ape* v5.0 (Paradis & Schliep 2019).

The priors on our parameters were: $\alpha \sim \text{Half-Normal}(0, 10)$, $\phi \sim \text{Exponential}(\mathbb{V}[\pi])$, $\gamma \sim \mathcal{N}(\text{logit}(\mathbb{E}[\pi]), 1)$, and $\beta \sim \mathcal{N}(0, 10)$, where $\mathbb{V}[\pi]$ and $\mathbb{E}[\pi]$ denote the empirical variance and mean, respectively, of observed $\pi$ across species. Each model contained one focal predictor of interest, the phylogenetic variance-covariance matrix, and four nuisance predictors (number of individuals in the species dataset; the mean raw read count; the mean read length; and the mean locus depth).

For each model, we included all species that were scored for the focal trait of interest (excluding species with missing data or for which the trait was not applicable). We ran each MCMC chain for 20,000 iterations, discarding the first half of the iterations as burn-in and thinning the remaining samples by retaining every 40th iteration. We assessed model convergence by examining parameter trace plots, marginal distributions, and joint marginal distributions. For each predictor discussed in the text, we report the 95% equal-tailed credible interval from the thinned, burned-in marginal distribution.

We assessed the significance of each parameter by determining whether the 95% equal-tailed credible interval contained the value 0; if it did, the parameter was deemed not significant. To compare effect sizes between predictors, we standardized them by multiplying the estimated effect size by the standard deviation of the predictor. In order to evaluate model adequacy (i.e., the ability of our model to describe the data), we used posterior predictive sampling (Gelman et al. 1996). For each model run, we randomly sampled 500 MCMC iterations, with replacement, from the burned-in, thinned, posterior distribution. For each sampled iteration, we recorded the estimated parameter values for the shape parameters of the Beta distribution used to model our response variable $a$ and $b$. We then simulated draws from a Beta distribution parameterized by $a_i$ and $b_i$ for sampled iteration $i$. Those posterior predicted values of $\pi$ were compared to the observed value of $\pi$ for each species to determine whether the model as a whole was effectively capturing patterns in the data, as well as to determine whether diversity for specific species was well or poorly predicted by our model (Figure S9).

Because range extent and number of ecoregions were both significant predictors of genetic diversity (Figure 2), but were also highly correlated (Figure S4), we ran an additional analysis to try to get closer to ascertaining causality. Number of ecoregions is necessarily correlated with range extent, so we were concerned that the significance of ecoregions was being driven by its correlation with range extent. To test whether this suspicion was correct, we analyzed diversity as a function of number of ecoregions *per* unit range extent. The mean estimated effect size of this hybrid predictor was -0.003, with a 95% equal-tailed posterior credible interval of -0.0072 – 0.0009 which was not significantly different from zero. As a result, we cannot conclusively rule out the possibility that the effect of ecoregions on diversity is entirely being driven by the correlation of ecoregions with range extent, and we thus drop ecoregions from further investigation.

To assess the explanatory power of each model, we calculate a Bayesian $R^2$ following the recommendations of Gelman et al. (2019). Specifically, we work in the 'link' space of the model and, for each sampled iteration in the MCMC, we calculate the ratio of the variance in the fitted means to the sum of the variance of fitted means and residual means. For a given iteration $s$, this is as follows:

$$\mathbb{E}[\theta_s] = \gamma_s + \sum_M \beta_{sm} \vec{X}_m \tag{6}$$

$$\mathbb{V}_{\text{fit}} = \text{Var}\left(\mathbb{E}[\theta_s]\right) \tag{7}$$

$$\mathbb{V}_{\text{resid}} = \text{Var}\left(\theta_s - \mathbb{E}[\theta_s]\right) \tag{8}$$

$$R_s^2 = \frac{\mathbb{V}_{\text{fit}}}{\mathbb{V}_{\text{fit}} + \mathbb{V}_{\text{resid}}} \tag{9}$$

**Table S1:** Population genetic theory offers several hypotheses about factors that govern levels of genetic diversity within and among species. Here we focus on population size, connectivity, and demography. These are difficult to quantify directly, so we present a suite of biotic and abiotic traits that have been hypothesized to predict species-level genetic diversity given their relationships to population demography. We include citations to support these predicted linkages from theory and empirical examples where possible. Arrows indicate the expected direction of association with species-level genetic diversity. Note that some of the below traits, including all dispersal-related traits, have predictions for both increasing and decreasing species-level genetic diversity depending on the processes at play.

| **Population process shaping patterns of diversity** | **Abiotic and biotic traits used as proxies for column 1** | **Predicted correlation of proxy with genetic diversity** | **Rationale for predicted effect of abiotic and biotic traits on genetic diversity (via their relationships to factors in column 1)** | **Supporting theory (specific to trait listed)** | **Empirical examples linking trait to genetic diversity** |
|---|---|---|---|---|---|
| **Census population size ↑** | Adult body size | ↓ | Species with smaller body sizes tend to have larger census population sizes and therefore have larger $N_e$, increasing genetic diversity. | Brown 1995; White et al. 2007 | *Matches prediction:* Buffalo 2021; Dalongeville et al. 2016*; De Kort et al. 2021*; Eo et al. 2011; Leffler et al. 2012; Lewin & Eyre-Walker 2026; Mackintosh et al. 2019; Mitton & Lewis Jr 1989; Romiguier et al. 2014; Waples 1991 *No relationship:* Doyle et al. 2015; Pinsky & Palumbi 2014* *Opposite of prediction:* Donati et al. 2021 |

*Continued on next page*

*Supplemental Table 1, continued*

| Population process shaping patterns of diversity | Abiotic and biotic traits used as proxies for column 1 | Predicted correlation of proxy with genetic diversity | Rationale for predicted effect of abiotic and biotic traits on genetic diversity (via their relationships to factors in column 1) | Supporting theory (specific to trait listed) | Empirical examples linking trait to genetic diversity |
|---|---|---|---|---|---|
| | Egg size | ↓ | Larger egg size predicts lower fecundity: lower fecundity is correlated with smaller population sizes, and lower genetic diversity. | Kimura & Crow 1963, eq. 18 | *Matches prediction:* Lewin & Eyre-Walker 2026; Mitton & Lewis Jr 1989; Romiguier et al. 2014 (propagule size and fecundity were strongest predictors in study); *No relationship:* Barry et al. 2022*; Mackintosh et al. 2019 *Opposite of prediction:* Martinez et al. 2018* Waples 1991 (for marine fishes); |
| | Spawning mode (trait "increases" from internally fertilized to nesting to broadcasting) | ↑ | Expect free-floating, broadcast modes of spawning to have greatest fecundity and thus larger population sizes (inverse of egg size), and higher genetic diversity. | | *Matches prediction:* Barry et al. 2022* *No relationship:* Cahill et al. 2017 |
| | Range extent | ↑ | Species with larger geographic ranges will have larger census population sizes and therefore have larger $N_e$, increasing genetic diversity. | Wilkins 2004 | *Matches prediction:* Hamrick & Godt 1996; James & Eyre-Walker 2020; Leffler et al. 2012* *Weak support for prediction:* De Kort et al. 2021* *No relationship:* Doyle et al. 2015; Lawrence & Fraser 2020*; Mackintosh et al. 2019 |

*Continued on next page*

*Supplemental Table 1, continued*

| **Population process shaping patterns of diversity** | **Abiotic and biotic traits used as proxies for column 1** | **Predicted correlation of proxy with genetic diversity** | **Rationale for predicted effect of abiotic and biotic traits on genetic diversity (via their relationships to factors in column 1)** | **Supporting theory (specific to trait listed)** | **Empirical examples linking trait to genetic diversity** |
|---|---|---|---|---|---|
| **Demographic fluctuations & Bottlenecks** | Mean range latitude | ↓ | Latitudes further away from the equator are predicted to experience less abiotic stability, which may lead to more bottlenecks (less demographic stability), decreasing genetic diversity; similar arguments to latitudinal species diversity gradients: expect greater abundance, more productivity, etc. in tropics. | Lawrence & Fraser 2020 (presents multiple predictions) | *Matches prediction:* Adams & Hadly 2013*; Manel et al. 2020; Miraldo et al. 2016* *Mixed support for prediction:* Clark & Pinsky 2024*; De Kort et al. 2021* *No relationship:* Pinsky & Palumbi 2014* *Opposite of prediction:* Figuerola-Ferrando et al. 2023* |
| | Egg size | ↓ and ↑ | Larger egg size predicts lower fecundity (K-strategists), which may lead to slower recovery from bottlenecks, decreasing genetic diversity (compared to r-strategists). Greater reproductive variance associated with smaller eggs leads to decreased genetic diversity. | Romiguier et al. 2014 – Supplemental Information (↓ prediction) Crow & Kimura 1970; Hedgecock & Pudovkin 2011 (↑ prediction) | *Matches ↓ prediction:* Lewin & Eyre-Walker 2026; Romiguier et al. 2014 *Matches ↑ prediction:* Martinez et al. 2018 (note maximum fecundity is used rather than egg size) |
| | Spawning | ↓ | Greater reproductive variance associated with broadcast spawning leads to decreased genetic diversity. | Crow & Kimura 1970; Hedgecock & Pudovkin 2011 | *No relationship:* Cahill et al. 2017 |

*Continued on next page*

*Supplemental Table 1, continued*

| **Population process shaping patterns of diversity** | **Abiotic and biotic traits used as proxies for column 1** | **Predicted correlation of proxy with genetic diversity** | **Rationale for predicted effect of abiotic and biotic traits on genetic diversity (via their relationships to factors in column 1)** | **Supporting theory (specific to trait listed)** | **Empirical examples linking trait to genetic diversity** |
|---|---|---|---|---|---|
| | Iteroparity | ↑ and ↓ | Iteroparity allows for multiple matings in a lifetime, which may reduce reproductive variance generated by lucky/unlucky years (compared to the single reproductive event of semelparous species), increasing genetic diversity. However, variance in longevity may increase lifetime reproductive variance across continually reproducing individuals, decreasing genetic diversity. Additionally, iteroparous species often have lower fecundity (and age-specific mortality), which may lead to decreased diversity. | Waples 2006 (↑ prediction)<br>Barry et al. 2022 (↓ prediction) | *Weak support for ↑ prediction:* Dalongeville et al. 2016*<br>*No support:* Hamrick & Godt 1996 |
| | Adult body size | ↓ | Large-bodied species are often long-lived. If they have a Type III life history curve (high juvenile mortality and low adult mortality), which is typical of many marine fishes and invertebrates, as well as sea turtles, they may have greater variance in lifetime reproductive output, reducing genetic diversity. | Barry et al. 2022; Waples 2016 | *Matches prediction:* Barry et al. 2022* |

*Continued on next page*

*Supplemental Table 1, continued*

| **Population process shaping patterns of diversity** | **Abiotic and biotic traits used as proxies for column 1** | **Predicted correlation of proxy with genetic diversity** | **Rationale for predicted effect of abiotic and biotic traits on genetic diversity (via their relationships to factors in column 1)** | **Supporting theory (specific to trait listed)** | **Empirical examples linking trait to genetic diversity** |
|---|---|---|---|---|---|
| **Population connectivity** Dispersal ↓ and ↑ | Egg size | ↓ and ↑ | Larger egg size is correlated with less dispersal, less dispersal facilitates differentiation among demes, increasing species-level genetic diversity. Low dispersal also inhibits demes rescue after local bottlenecks and can lead to the loss of sink populations, both of which decrease genetic diversity at the species level. | Villarino et al. 2018 (smaller eggs should be carried further on currents and resist sinking) | (Literature exists relating egg size to population structure, e.g., Riginos et al. 2011, but not examples relating to species-level genetic diversity) |
| | Planktonicity | ↓ and ↑ | Presence of a free-floating life stage leads to increased dispersal, homogenizing genetic diversity among demes, and lowering species-level genetic diversity. High dispersal also helps rescue demes from local bottlenecks and maintains sink populations, both of which increase genetic diversity at the species level. | | *Moderate support for ↓ prediction:* Hamrick & Godt 1996 |
| | Spawning mode (trait "increases" from internally fertilized to nesting to broadcasting) | ↓ and ↑ | Expect free-floating broadcast modes to have greater (progeny/parent) dispersal, homogenizing genetic diversity among demes, and lowering species-level genetic diversity. High dispersal also helps rescue demes from local bottlenecks and maintains sink populations, both of which increase genetic diversity at the species level. | | *No relationship:* Cahill et al. 2017 |

*Continued on next page*

*Supplemental Table 1, continued*

| **Population process shaping patterns of diversity** | **Abiotic and biotic traits used as proxies for column 1** | **Predicted correlation of proxy with genetic diversity** | **Rationale for predicted effect of abiotic and biotic traits on genetic diversity (via their relationships to factors in column 1)** | **Supporting theory (specific to trait listed)** | **Empirical examples linking trait to genetic diversity** |
|---|---|---|---|---|---|
| | Pelagic larval duration (PLD) | ↓ and ↑ | Longer pelagic larval duration leads to increased (progeny/parent) dispersal, homogenizing genetic diversity among demes, and lowering species-level genetic diversity. High dispersal also helps rescue demes from local bottlenecks and maintains sink populations, both of which increase genetic diversity at the species level. | | *No relationship:* Donati et al. 2021 (There is extensive literature relating PLD to population structure but not examples relating to species-level genetic diversity) |
| | Planktotrophy | ↓ and ↑ | Increased planktotrophy leads to increased (progeny/parent) dispersal, homogenizing genetic diversity among demes, and lowering species-level genetic diversity. High dispersal also helps rescue demes from local bottlenecks and maintains sink populations, both of which increase genetic diversity at the species level. | | *No relationship:* Cahill et al. 2017 |
| | Philopatry | ↓ and ↑ | Adult philopatry reduces average parent-offspring distance; less dispersal facilitates differentiation among demes, increasing species-level genetic diversity. Low dispersal also inhibits demes rescue after local bottlenecks and can lead to the loss of sink populations, both of which decrease genetic diversity at the species level. | | (There is extensive literature relating philopatry to population structure but not examples relating to species-level genetic diversity) |

*Continued on next page*

*Supplemental Table 1, continued*

| **Population process shaping patterns of diversity** | **Abiotic and biotic traits used as proxies for column 1** | **Predicted correlation of proxy with genetic diversity** | **Rationale for predicted effect of abiotic and biotic traits on genetic diversity (via their relationships to factors in column 1)** | **Supporting theory (specific to trait listed)** | **Empirical examples linking trait to genetic diversity** |
|---|---|---|---|---|---|
| | Benthicity | ↓ and ↑ | Benthicity reflects adult dispersion ability via the degree to which they are associated with the ocean floor: more benthic species maintain a stronger connection to the bottom, which predicts less dispersal, and higher genetic diversity. Low dispersal also inhibits demes rescue after local bottlenecks and can lead to the loss of sink populations, both of which decrease genetic diversity at the species level. | | *Matches ↑ prediction:* Romiguier et al. 2014<br>*Mixed support for ↑ prediction:* Dalongeville et al. 2016* |
| **Population connectivity**<br>Range size and complexity ↑ | Range extent | ↑ | Larger ranges lead to greater geographic distances that can separate individuals, increasing the waiting time to coalescence, and increasing genetic diversity. | Maruyama 1972; Wilkins 2004 | *Matches prediction*: Leffler et al. 2012<br>*No relationship*: Lewin & Eyre-Walker 2026 |
| | Number of ecoregions (that the species' range contains) | ↑ | More ecoregions implies ecological discontinuities, which may act as direct (dispersal) or indirect (selection) barriers to dispersal, increasing the time to coalescence (more environmental distance over which isolation can happen), and increasing genetic diversity. | | |

*Authors use within-deme genetic diversity, not species-level diversity.

**Table S2**. Biotic and abiotic predictors of species-level genetic diversity included in analysis. "Trait values" describes the range of possible values species could have for each predictor. "Significant" describes whether the effect size of the predictor was inferred to be significantly different from zero. The column "n" gives the sample size (uncorrected for phylogeny) for each analysis (i.e., the number of species included, of the total 91 species included in this study). The final column gives the mean (unstandardized) effect size of the predictor, as well as the lower and upper bounds of the 95% equal-tailed credible interval of the marginal distribution of the effect size.

| **Predictor** | **Trait values** | **n** | **Significant** | **Mean effect (95% CI)** |
|---|---|---|---|---|
| Benthicity | absent/intermediate/present | 91 | no | -5.20e-02 (-4.10e-01, 2.60e-01) |
| Body size | continuous | 91 | no | -5.40e-02 (-1.50e-01, 6.70e-02) |
| Ecoregions | integer | 91 | yes | 6.00e-03 (1.10e-05, 1.20e-02) |
| Egg size | continuous | 66 | no | -3.30e-03 (-1.50e-02, 8.10e-03) |
| Iteroparity | absent/present | 91 | no | -3.80e-01 (-9.70e-01, 1.80e-01) |
| Latitude | continuous | 91 | no | -3.00e-03 (-1.20e-02, 6.80e-03) |
| PLD | continuous | 85 | no | -7.30e-04 (-4.70e-03, 3.20e-03) |
| Philopatry | absent/present | 87 | no | -4.20e-01 (-9.10e-01, 5.00e-02) |
| Planktonicity | absent/present | 91 | yes | 5.80e-01 (3.20e-02, 1.20e+00) |
| Planktotrophy | absent/intermediate/present | 91 | no | 1.70e-01 (-8.90e-02, 4.20e-01) |
| Spawning | absent/intermediate/present | 91 | no | 1.10e-01 (-7.00e-02, 3.10e-01) |
| Range extent | continuous | 91 | yes | 5.20e-01 (1.60e-02, 9.60e-01) |

**Table S3**. Estimated effect sizes for each "nuisance" variable included as a covariate in phylogenetic beta regression analyses of species-level genetic diversity. As described in the *Methods*, the effect of each ecological predictor was tested in a model that included that primary predictor of interest (rows in below table) and four nuisance variables (columns in below table). Each cell below gives the mean and 95% equal-tailed credible interval for the estimated effect size of each nuisance variable in each of the 12 models built, one model for each ecological predictor. Effect sizes for which the 95% CI does not contain 0 are written in bold.

| | Number of samples | Mean Raw Read Counts | Mean Read Length | Mean Locus Depth |
|---|---|---|---|---|
| latitude | −0.001 [−0.003, 0.001] | **−0.640 [−1.049, −0.251]** | **−0.006 [−0.011, −0.000]** | **0.006 [0.003, 0.009]** |
| ecoregions | −0.001 [−0.003, 0.000] | **−0.609 [−0.989, −0.208]** | **−0.006 [−0.011, −0.001]** | **0.005 [0.003, 0.008]** |
| range extent | −0.001 [−0.003, 0.001] | **−0.624 [−1.006, −0.195]** | **−0.006 [−0.011, −0.001]** | **0.006 [0.003, 0.009]** |
| body size | −0.001 [−0.003, 0.000] | **−0.648 [−1.079, −0.259]** | −0.005 [−0.010, 0.000] | **0.006 [0.003, 0.009]** |
| egg size | −0.001 [−0.002, 0.000] | **−0.891 [−1.274, −0.504]** | −0.002 [−0.008, 0.005] | 0.002 [−0.002, 0.006] |
| iteroparity | −0.001 [−0.003, 0.000] | **−0.662 [−1.111, −0.282]** | −0.005 [−0.010, 0.000] | **0.005 [0.002, 0.009]** |
| philopatry | −0.001 [−0.003, 0.001] | **−0.701 [−1.145, −0.267]** | **−0.006 [−0.011, −0.000]** | **0.007 [0.003, 0.010]** |
| spawning | −0.002 [−0.004, 0.000] | **−0.714 [−1.175, −0.253]** | **−0.006 [−0.011, −0.000]** | **0.006 [0.003, 0.009]** |
| planktotrophy | −0.001 [−0.003, 0.000] | **−0.651 [−1.066, −0.253]** | **−0.005 [−0.010, −0.000]** | **0.006 [0.002, 0.009]** |
| PLD | −0.001 [−0.003, 0.001] | **−0.637 [−1.107, −0.203]** | −0.005 [−0.010, 0.000] | **0.006 [0.003, 0.009]** |
| planktonicity | −0.002 [−0.004, 0.000] | **−0.701 [−1.208, −0.247]** | **−0.006 [−0.011, −0.000]** | **0.006 [0.003, 0.009]** |
| benthicity | −0.001 [−0.003, 0.000] | **−0.683 [−1.041, −0.268]** | −0.005 [−0.010, 0.001] | **0.005 [0.002, 0.008]** |

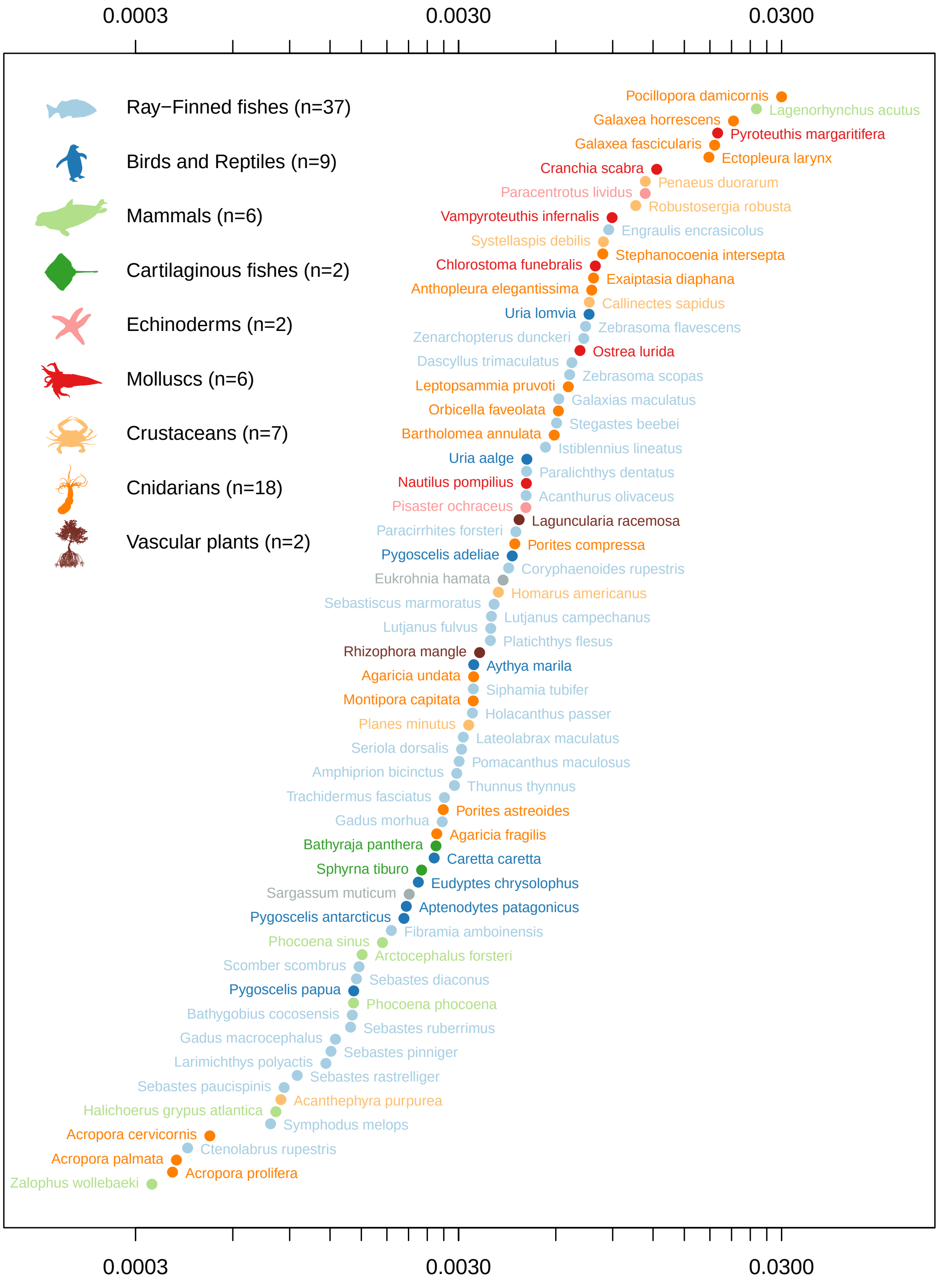


**Figure S1**. Species-level genetic diversity estimated for each species included in analysis. Species diversity estimates are colored by broad taxonomic group and sorted by value.

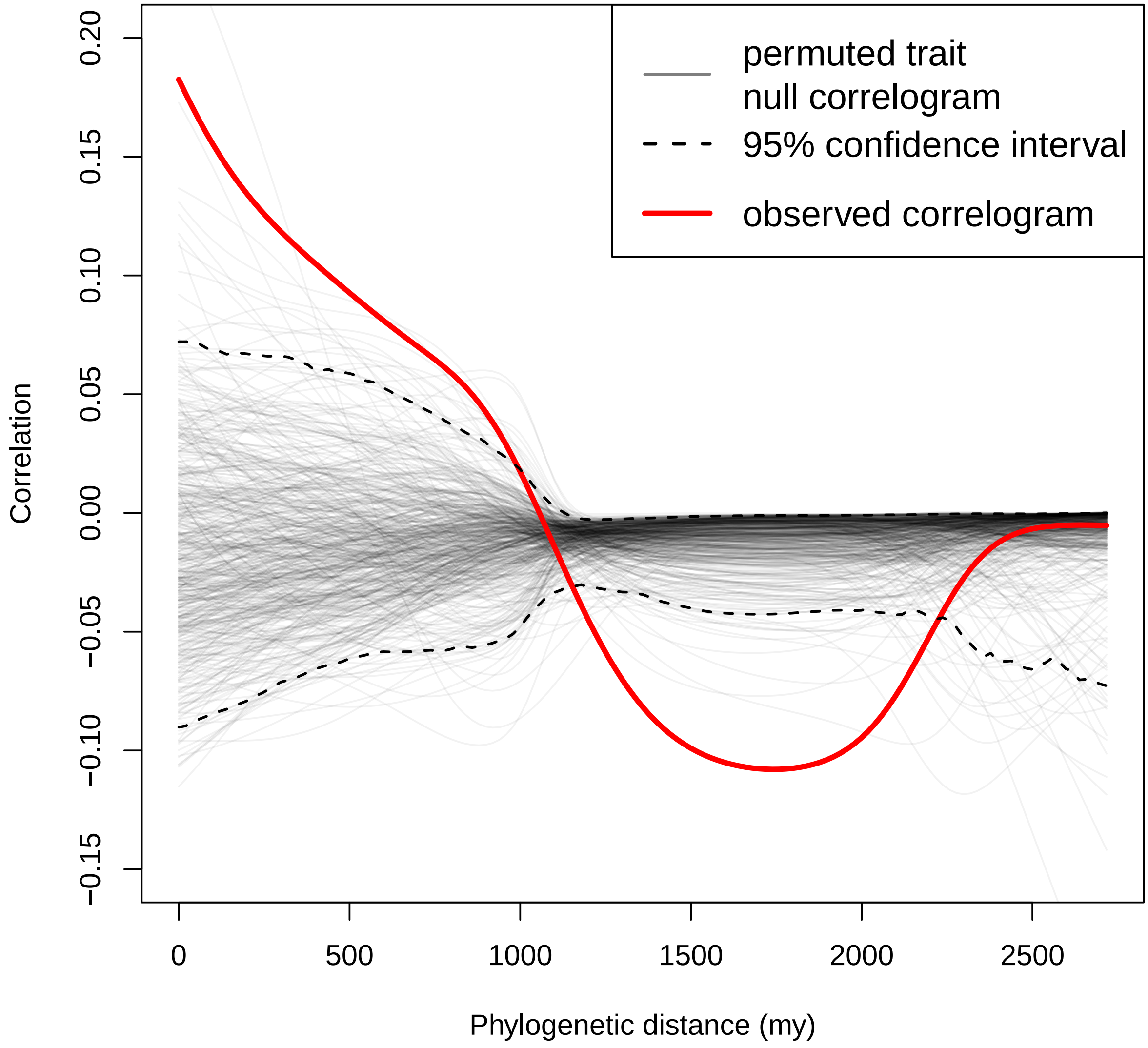


**Figure S2**. Phylogenetic correlogram showing the estimated correlation between phylogenetic relatedness and species-level genetic diversity at different temporal depths.

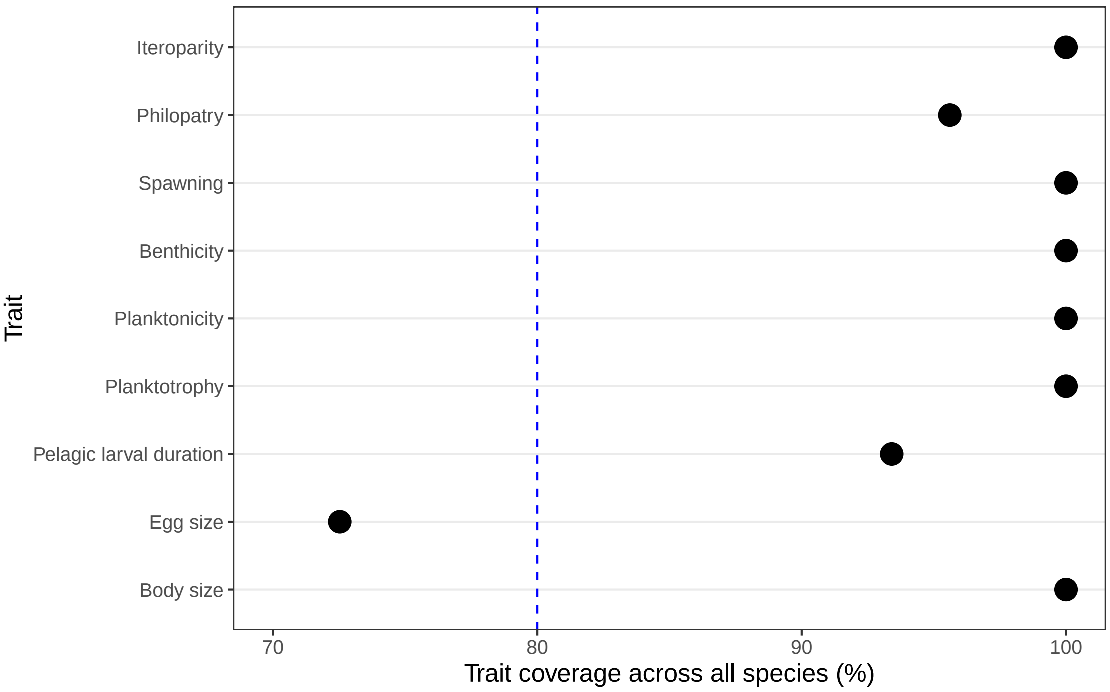


**Figure S3**. Trait coverage for each of the nine biotic traits included in this study; coverage calculated as a percentage across all 91 species in the study. Note the *x*-axis starts at 70% (rather than 0). The vertical blue dashed line denotes 80%, a commonly accepted conservative threshold for trait coverage. (Note the three abiotic predictors were available for all datasets in this study.)

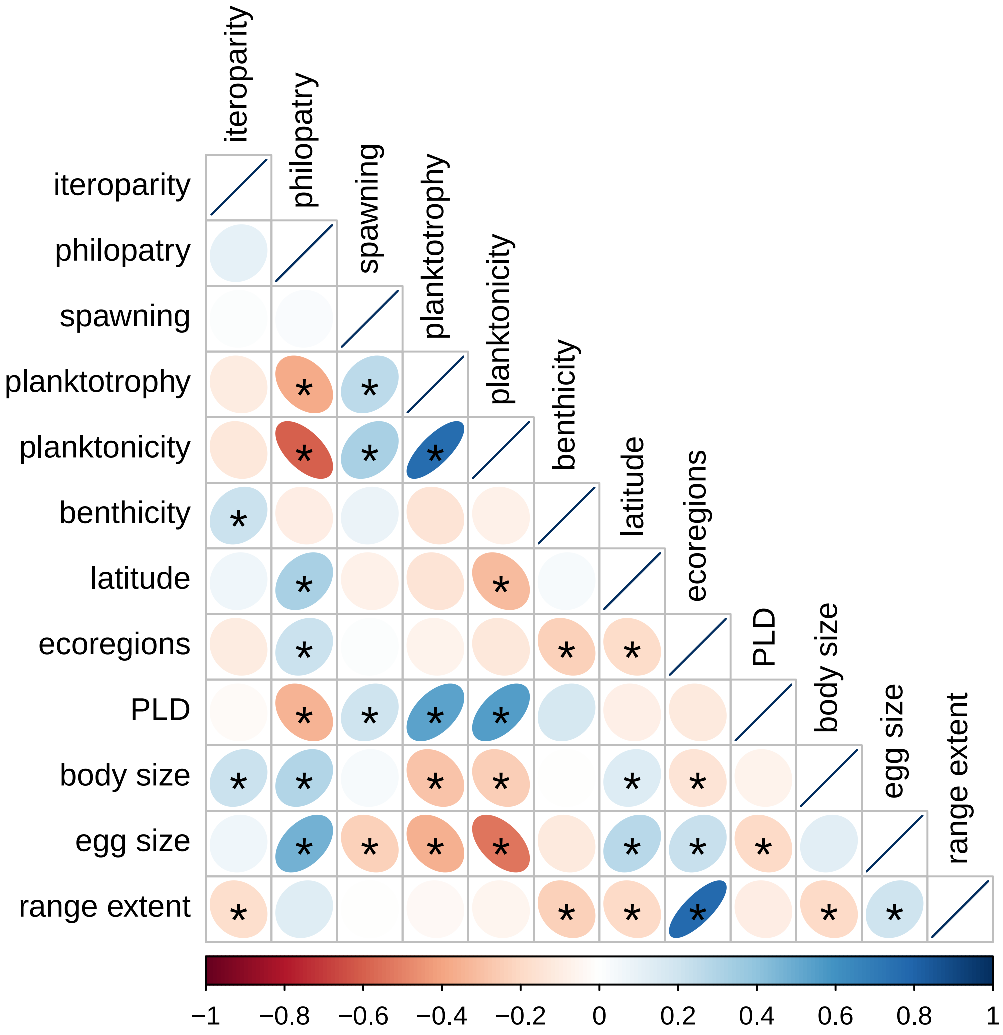


**Figure S4**. Kendall rank correlations (using pairwise complete observations) between all biotic and abiotic predictors analyzed. Blue and red colors indicate positive and negative correlations, respectively, and color intensity is proportional to the absolute value of the correlation. Significant correlations (assessed using the R function `cor.test` in the `stats` package using a two-sided alternative and a 95% confidence level) are indicated with an asterisk.

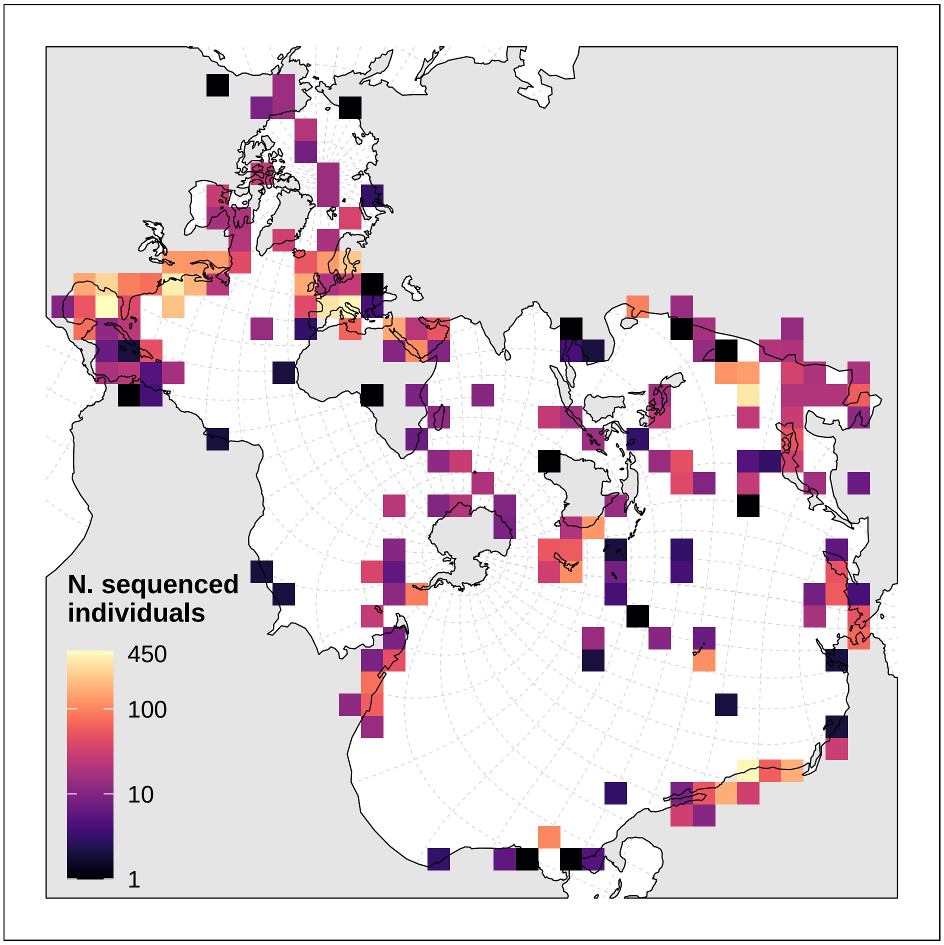


**Figure S5**. Spatial distribution and sampling intensity of sequenced individual marine organisms included in this study ($N$ = 9,325 total). Colors represent the number of sequenced individuals within each grid square. Map is in the Spilhaus projection.

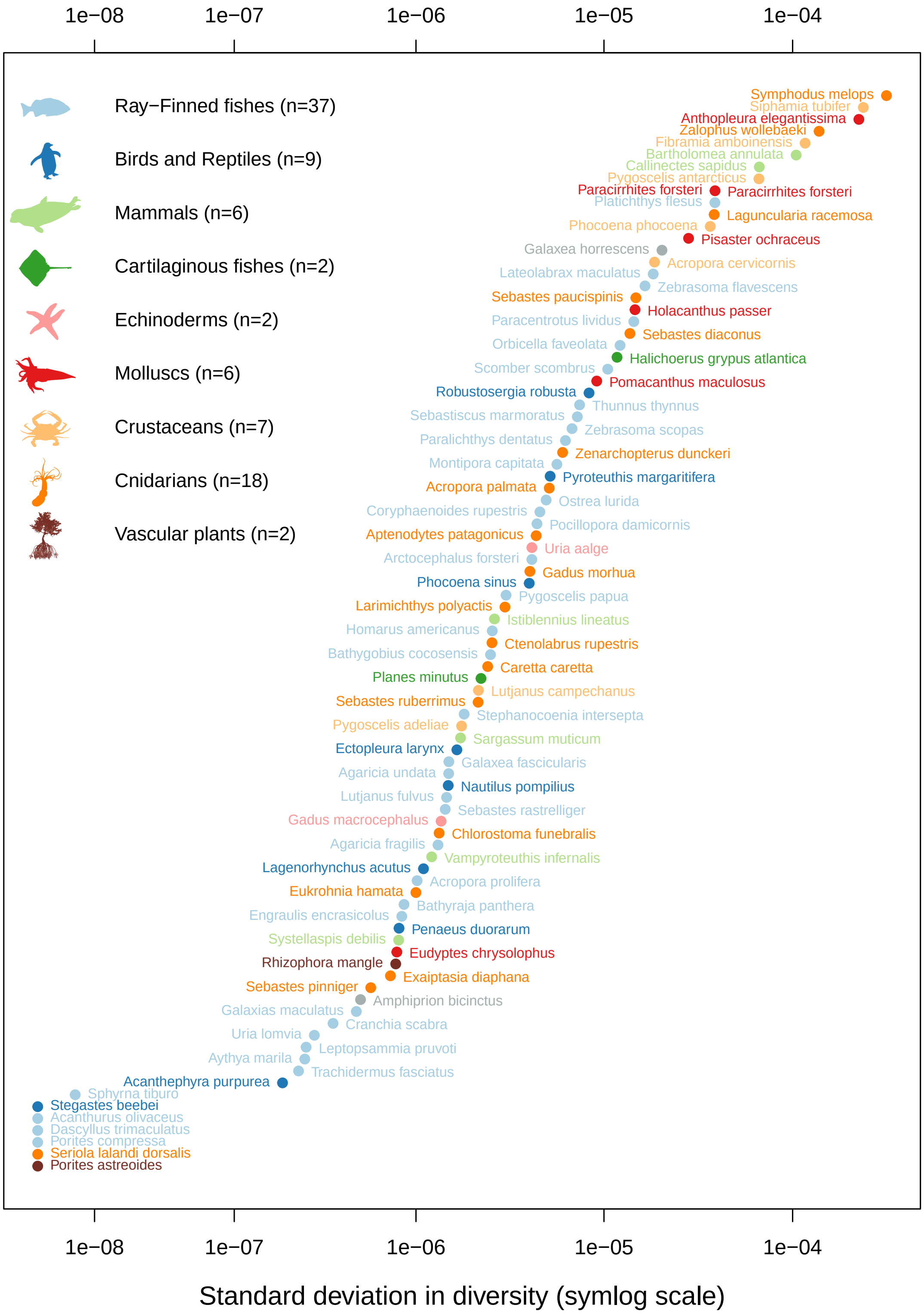


**Figure S6**. Standard deviation of diversity estimates across unique locations for each species included in analysis. The standard deviations of the species diversity estimates are colored by broad taxonomic group and sorted by value. Values are plotted on a symlog scale.

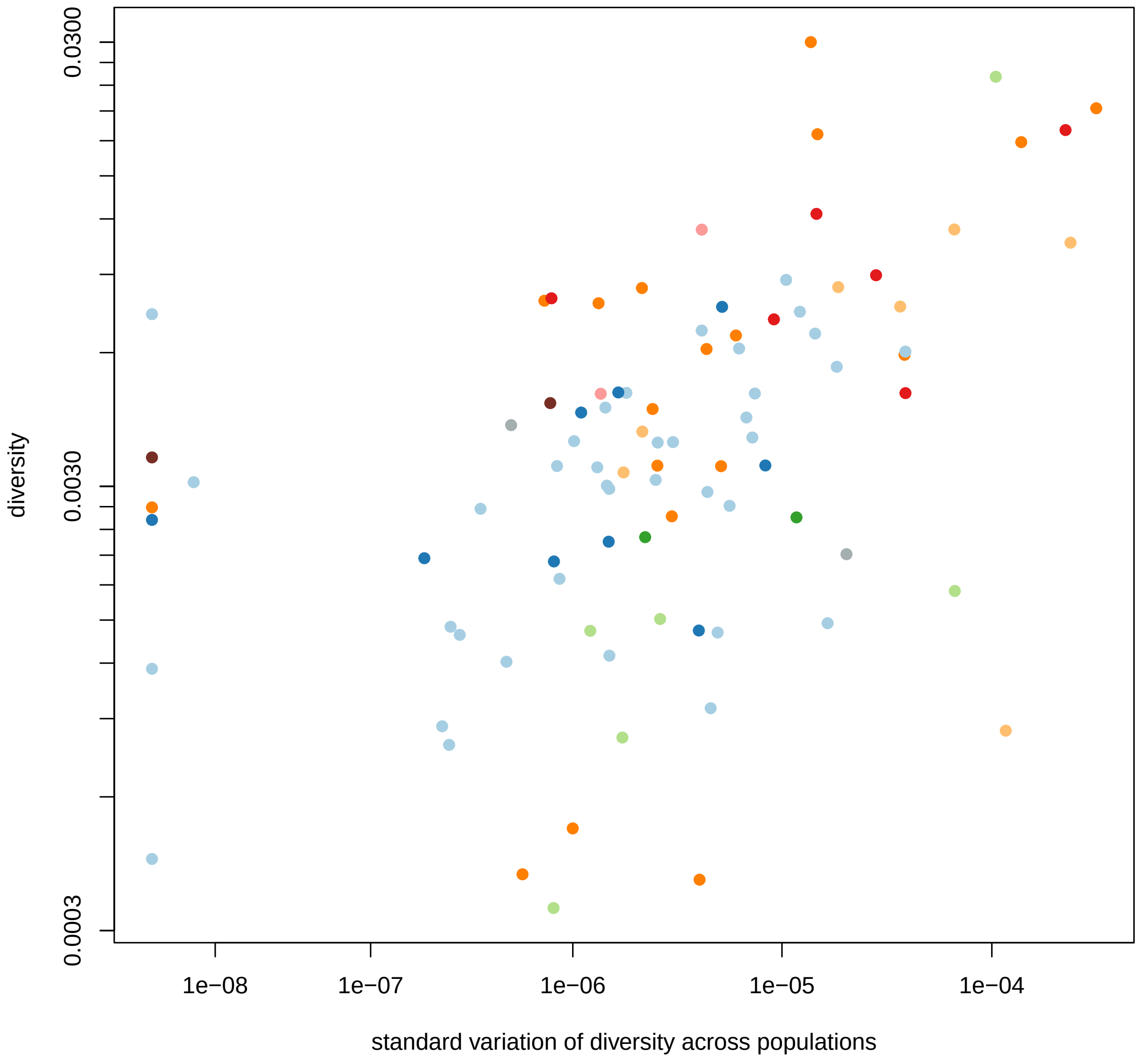


**Figure S7**. Standard deviation of diversity estimates across unique locations for each species (symlog scale) plotted against the total diversity estimate for that species (log scale). Points colored by broad taxonomic group.

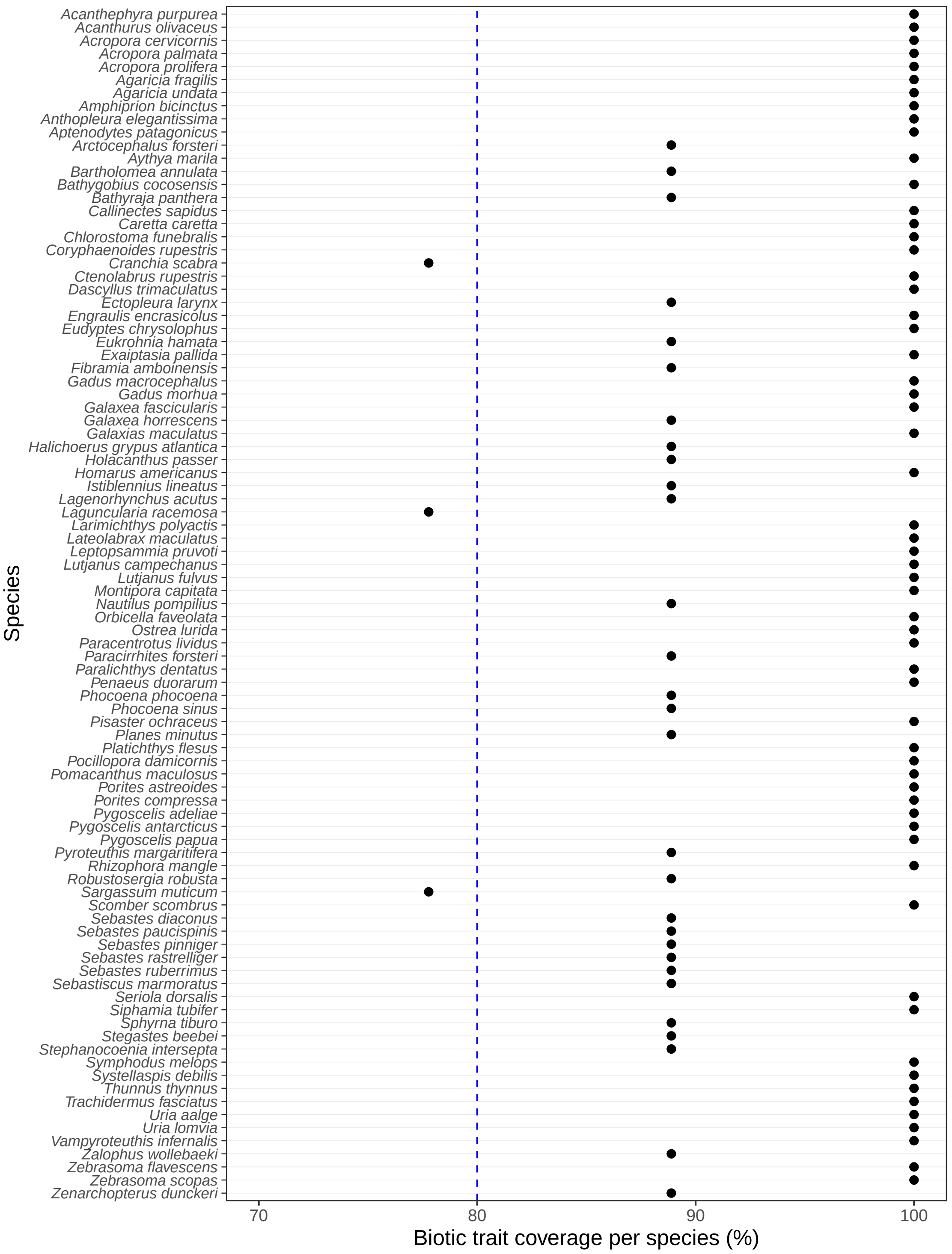


**Figure S8**. Biotic trait coverage for each species in the study ($N$ = 91) as a percentage. There are nine biotic traits total. Note the $x$-axis starts at 70% (rather than 0). Species are ordered alphabetically by Latin name. The vertical blue dashed line denotes 80%, a commonly accepted conservative threshold for trait coverage.

### Appendix S1: Extended definitions and coding of biotic traits

This appendix contains detailed explanations of how we defined the nine biotic traits included in this study. Headers indicate the variable name used in the paper followed parenthetically by the variable name used in archived data files and scripts. Bullet points denote possible values for discrete traits; values after the colon specify the short-hand coding (capital letters) used to record trait data, numerical values represent the dummy numerical values that characters were transformed into for modeling, and bracketed text specifies how discrete traits are shown in Figure 1. See *Supplemental Methods – Collecting biotic, abiotic, and phylogenetic predictors* for a description of how trait values were obtained.

---

***Iteroparity ("Generational_Structure")***

This variable captures variability in the number of times an individual can breed in its lifetime. Iteroparous species can breed more than once in their lifetime, whereas semelparous species only breed once.

- Iteroparous: I = 1 [present]
- Semelparous: S = 0 [absent]

***Philopatry ("ReturnToSpawningGround")***

We defined philopatry as the intentional movement of reproductively mature individuals back to an established spawning or breeding area to reproduce. Species where evidence indicated such migration were coded as TRUE and species lacking such evidence were coded as FALSE. For species that could plausibly have such migratory life histories (i.e., *Bathyraja panthera* and *Pyroteuthis margaritifera*, with large bodied and strong swimming adults with poorly described life histories with related species known to be philopatric), we scored this trait as unknown.

- Philopatric: TRUE = 1 [present]
- Not philopatric: FALSE = 0 [absent]
- Unknown trait state: NA [missing data]

***Spawning ("Spawning_mode")***

Marine taxa have a wide array of reproductive strategies with respect to fertilization and subsequent embryonic development and parental care. Our primary focus here was on capturing variation in how far embryos disperse from parents. For animals, eggs or developing young can be internally fertilized and then retained in a parents' body or on a parent's body in specialized brooding structures (pouches, chambers, sacs). Alternatively, animals can lay eggs in nests or other benthically attached structures; parents often (but not always) then disassociate from the laid eggs. Finally, many fishes and invertebrates release free floating embryos, typically preceded by broadcast spawning of gametes. We coded all mammals as "I" and all birds as "N"; fishes and invertebrates had a diverse mix of values. For plants and algae, we applied similar logic as for animals and scored species that retain young on the parent as "I".

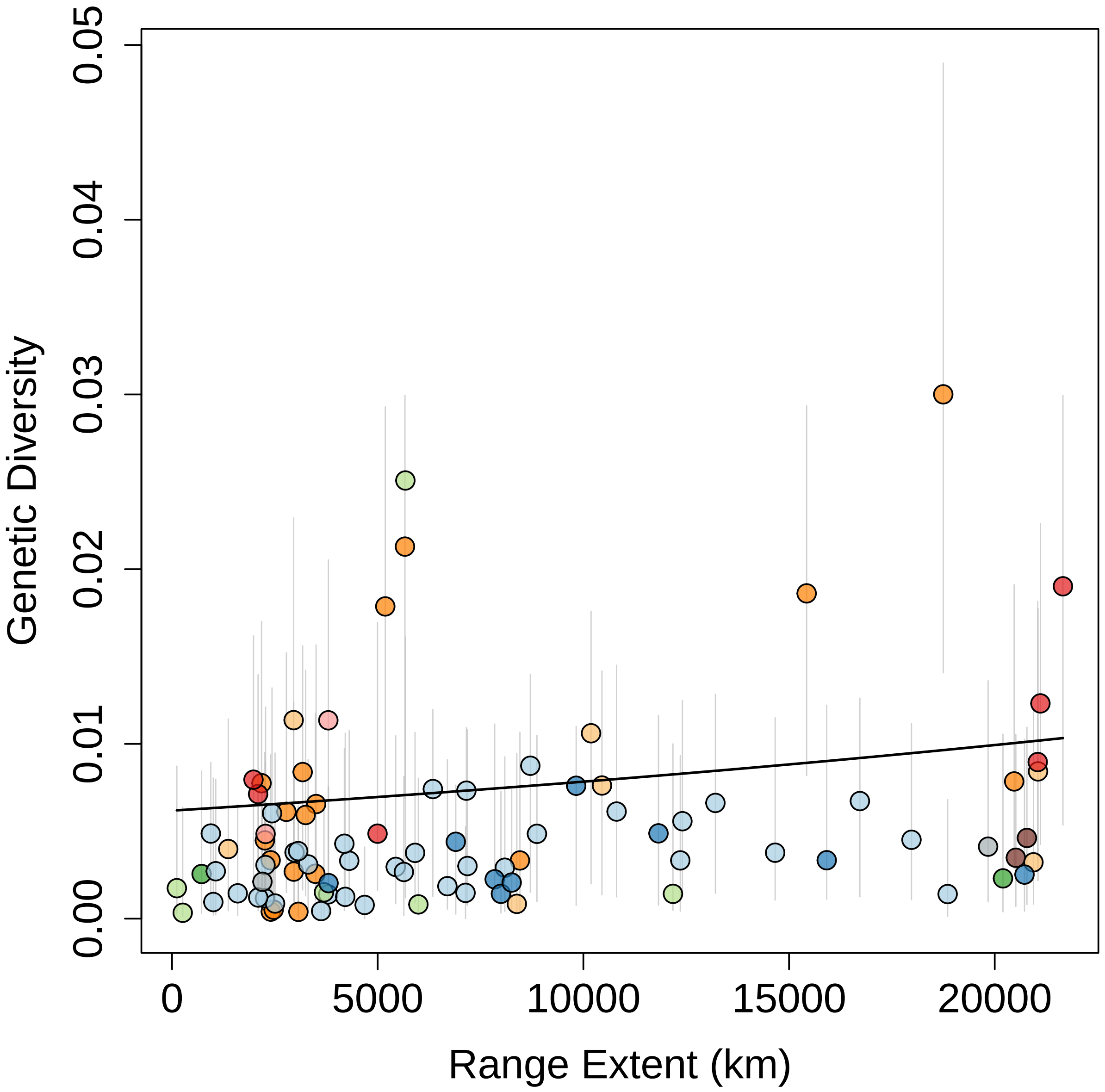


**Figure S9**. Relationship between range extent and species-level genetic diversity. Points are colored by broad taxonomic group as in Figure 1. Vertical gray lines show the 95% equal-tailed quantile of the posterior predictive distribution of estimates for diversity for each species.

- Embryos internally fertilized and attached to parents: I = 0 [absent]
- Embryos in a nest or otherwise benthically attached: N = 1 [intermediate]
- Embryos free floating: F = 2 [present]

***Benthicity ("isBenthic")***

This variable attempts to capture the diversity of relationships between organisms and a fixed location (e.g., for birds this can be nesting sites on land). We considered organisms to be "never" benthic if their entire life cycle was dissociated from any bottom surface (e.g., anchovy that have planktonic larvae and pelagic swimming adults or dolphins that give birth at sea to swimming young). "Sometimes" benthic organisms have a mixture of life stages that are benthic and pelagic, for example many fishes and invertebrates with planktonic larvae and benthic adult stages. "Always" benthic organisms live their entire life with a strong connection to the bottom (such as hydrozoans and anemonefishes that lack planktonic larvae).

- Always: A = 2 [present]
- Sometimes: S = 1 [intermediate]
- Never: N = 0 [absent]

***Planktonicity ("isPlanktonic_atanypoint")***

We defined planktonicity to capture whether or not species were planktonic – floating or drifting and not able to actively swim against ocean currents – at any point in their lifetime. This variable attempts to distinguish species that have at least one planktonic life stage (larval, adult, or both) from species that entirely lack a planktonic stage. For example, all mammals, birds and elasmobranchs were coded as FALSE.

- Species is planktonic at any point in their lifecycle: TRUE = 1 [present]
- Species is not planktonic at any point in their lifecycle: FALSE = 0 [absent]

***Planktotrophy ("Larval_feeding")***

Many marine species, especially invertebrates, have a larval stage that differs in morphology from their adult forms. One common way to classify differences in larval life history is based on larval feeding mode, which in turn influences dispersal (as well as development time and survival probabilities). We defined three categories of larval feeding for this trait: species that do not have planktonic larvae, species with non-feeding planktonic larvae, and species with feeding planktonic larvae. Species that do not have planktonic larvae produce progeny that develop directly into young that resemble miniature adults; these young do not spend time drifting or floating in the water column. Species with non-feeding planktonic larvae, often called lecithotrophic, produce energy-rich eggs that facilitate development from larvae into juveniles without the need to feed; as these larvae are planktonic, they drift in the water column while developing, but development is relatively quick given the well-provisioned eggs. Finally, feeding (or planktotrophic) planktonic larvae derive from eggs that provide only enough energy to produce a larvae; these larvae must then spend time drifting in the water column feeding on particulate food to fuel development into juveniles, which usually takes longer than in (well-provisioned) non-feeding planktonic larvae. All fish with planktonic larvae are feeding (coded "P"). For corals, we coded species that do not have zooxanthellae in their larvae and rely fully on maternal provisioning as "L", and coded corals with larvae that eat at some point and/or are autotrophic (contain zooxanthellae that can photosynthesize and provide nutrition "on the go") as "P".

- No planktonic larvae: N = 0 [absent]
- Non-feeding (lecithotrophic) planktonic larvae: L = 1 [intermediate]
- Feeding (planktotrophic) planktonic larvae: P = 2 [present]

***Pelagic larval duration ("PLD_point2")***

A continuous variable representing the mean or midpoint of the estimates available for each species for the length of time (in days) that marine larvae spend in the plankton. We assigned this variable a value of 0 for species that have live birth or brood their young that become benthic adults (isBenthic=Always) and for species who are never benthic (isBenthic=Never); thus pelagic larval duration and benthicity are not conflated in our analyses.

***Egg size ("Fecundity_EggSize")***

A continuous variable in millimeters, representing the largest dimension of the egg or juvenile at birth or hatching (the propagule size, in the sense of Romiguier et al. (2014)).

***Body size ("BodySize")***

A continuous variable in centimeters, representing the largest dimension of an average organism of each species at its full-grown adult size. As corals and seagrasses are colonial, the meaning of body size is unclear. While we acknowledge that genet size (the collective size of a genotype that might include many discrete ramets or colonies) would be most appropriate in terms of scaling to likely reproductive output, genet sizes are largely unknown and therefore we used ramet or colony size. In some instances only colony area was provided, so we assumed a circular shape to derive the diameter.

### Appendix S2: Search strategy used to locate relevant genomic datasets in the INSDC

This appendix details the search strategy that we used to locate relevant genomic datasets in the International Nucleotide Sequence Database Collaboration (INSDC). Note that we accessed the INSDC from the North American portal – the National Center for Biotechnology Information (NCBI). We list the general types of filters that we applied, in the order we applied them, and the number (and type) of records remaining after each filtering step in our pipeline. We also detail the exact NCBI variable names and values that we used in our search and filtering pipeline to locate genomic datasets. NCBI uses BioProjects to denote a study or dataset, BioSamples to refer to individual organisms sampled, and SRA sequence records to refer to the genetic sequence data derived from a BioSample.

After the final filtering step listed below – passing species names to the World Register of Marine Species – we identified and dropped BioProjects (datasets) with fewer than 10 BioSamples (individuals) and 3 collection locations using R. We then curated the remaining potential datasets by hand. We dropped known species hybrids, salmonids, freshwater/inland samples, samples from locations not part of the native species' range, and samples from ancient time. We also dropped studies that were focused on specific organelles (e.g., chloroplast) and quantitative genetic studies. Final sequence samples were included if they had collection latitude and longitude, the raw FASTA file downloaded without errors, the FASTA sequence files were at the level of sampled individual (not pooled sequencing), and were not outliers with respect to read count.

---

***Overview of filters applied and number of records retained after each filtering step:***

⇒ 184,155 total BioProjects in INSDC on 10/19/2020

*Search BioProject database for all BioProjects that are:*
Non-human, non-viral, non-metagenomic, and non-bacterial
⇒ 32162 BioProjects returned

*Remove BioProjects related to:*
Gene expression, RefSeq, targeted loci, metagenomic, metabarcoding, exome, microbial
⇒ 19,495 BioProjects retained

*Download metadata from the Sequence Read Archive (SRA) for all sequence records in each of the above BioProjects.*
⇒ 948,446 SRA sequence records returned

*Retain SRA records with library strategies:*
WGS, WCS, WGA, RAD-Seq, OTHER
(removing: AMPLICON, ATAC-seq, Bisulfite-Seq, ChIA-PET, ChIP-Seq, CLONE, CLONEEND, CTS, DNase-Hypersensitivity, EST, FAIRE-seq, Hi-C, FINISHING, FL-cDNA, MBD, MeDIP, miRNA-Seq, MNase-Seq, MRE-Seq, ncRNA-Seq, POOLCLONE, RIP-Seq, RNA-Seq, SELEX, Synthetic-Long-Read, Targeted-Capture, TetheredChromatinConformationCapture, Tn-Seq, WXS)
⇒ 523,537 SRA sequence records retained

*Merge retained SRA records to the BioProject records using PRJ BioProject accession numbers.*

*Remove BioProjects that did not return any SRA records matching the above specified library strategies.*

⇒ 7,314 BioProjects retained

*Use the NCBI common tree online tool to view all taxonomic IDs assigned to the current list of sequence records. Remove sequences with a taxonomic identification of:*

Synthetic, metagenome, unidentified, bacteria, environmental, viral

*Download metadata from the BioSample database associated with each of the above sequence records.*

⇒ 505,932 SRA sequences from
⇒ 432,400 BioSamples from
⇒ 6,755 BioProjects retained representing
⇒ 18,556 purported species

*Retain records with species names present in the World Register of Marine Species (WoRMS).*

⇒ 114,865 SRA sequences from
⇒ 100,118 BioSamples from
⇒ 1,336 BioProjects retained representing
⇒ 1,738 purported species

---

##### ***Exact variables and terms used to locate and filter INSDC datasets:***

The exact search terms and filters that we used to identify the INSDC datasets analyzed in this publication follow. We use "!=" below to denote "does not equal". Bold headers refer to the NCBI databases, quoted terms denote variables/fields of metadata defined by NCBI within said database, and text beneath quoted terms denotes controlled vocabulary values of the specified variable.

##### **Initial BioProject search:**

```
"scope multiisolate"[Filter] OR "scope multispecies"[Filter] OR
"scope other"[Filter] NOT "org human"[Filter] NOT "org archaea"[Filter]
NOT "org viruses"[Filter] NOT "org bacteria"[Filter] NOT "metagenome"[Filter]
NOT "targeted locus loci"[Filter] NOT "clone ends"[Filter]
NOT "metagenomic assembly"[Filter] NOT "transcriptome gene expression"[Filter]
NOT "proteome"[Filter] NOT "epigenomics"[Filter] NOT "exome"[Filter]
NOT "bioproject protein"[Filter] NOT "bioproject gds"[Filter]
NOT "phenotype genotype"[Filter] NOT "metagenome"[All Fields] NOT "map"[Filter]
```

##### **BioProject filters:**

```
"project_data_type" !=

RefSeq Transcriptome or Gene expression
Targeted loci cultured
RefSeq Genome
RefSeq Genome sequencing and assembly
RefSeq Other
RefSeq Targeted Locus (Loci)
```

```
"project_target_material" !=
```

```
Phenotype
Proteome
Purified Chromosome
Reagent
Transcriptome
Targeted loci environmental
```

```
"project_target_capture" !=

Clone Ends
Exome
```

```
"organism_name" does not contain:

metagenome
```

```
"project_title" does not contain:

metagenome, Metagenome, Metagenomic, metagenomic, metabarcoding
transcriptome, Transcriptome, Transcriptomic
RNA
16S, 16s, 18S, 18s
CRISPR
gene expression, Gene Expression, Gene expression"
viral
microbiome, Microbiome, microbial, microbiota
bacteria, Bacteria
methylation
Plasmodium falciparum
```

#### SRA filters:

```
"library_strat" =
RAD-Seq
WCS
WGA
WGS
OTHER
```

```
"scientific_name" !=
metagenome, Metagenome
```

The below taxonomic IDs represent synthetic DNA, environmental DNA, metagenomic DNA, and bacterial DNA that we identified using the NCBI phylogenetic common tree online tool.

```
"taxid" !=
2293429
1440148
32630
1427524
81077
32644
496923
77133
669196
2282120
668369
727
2590021
28448
```

```
272943
2021969
527802
477819
562
227
83333
316407
703612
224308
1229511
1126252
1126251
```

**Taxonomy filters:**

```
"division" !=
Environmental samples
Bacteria
Viruses
```

**BioSample filters:**

```
"organism" =
# species returned by
worms::wormsbynames(match = FALSE, marine_only = TRUE)
```

We filtered remaining records to retain BioProjects with at least 10 BioSample IDs and at least three unique georeferenced sampling locations per species/BioProject combination (dataset). Finally, we manually dropped samples (or entire datasets) that were located in non-native areas of the species' ranges, were sampled from captive or lab individuals, and those derived from quantitative genetic studies, ancient time, or pooled genetic sequences. We also dropped known species hybrids, salmonids, and samples collected from freshwater or inland locations.

### Appendix S3: Citations for genomic datasets

This appendix contains citations for the scientific papers associated with the genetic sequence data utilized in this study. The BioProject accession number where the genetic data are archived in the National Center for Biotechnology Information (NCBI) is also included with each citation. Note some papers include multiple BioProjects and some genomic datasets (BioProjects) have multiple paper citations.

[7] Pim Bongaerts, Cynthia Riginos, Ramona Brunner, Norbert Englebert, Struan R Smith, and Ove Hoegh-Guldberg. Deep reefs are not universal refuges: reseeding potential varies among

coral species. *Science Advances*, 3(2):e1602373, 2017. *Genetic sequence data archived under NCBI BioProject Accession*: PRJNA361144.

[8] Elisa Boscari, Marco Abbiati, Fabio Badalamenti, Giorgio Bavestrello, Lisandro Benedetti-Cecchi, Rita Cannas, Angelo Cau, Carlo Cerrano, Giovanni Chimienti, Federica Costantini, Simonetta Fraschetti, Gianmarco Ingrosso, Ilaria A. M. Marino, Francesco Mastrototaro, Chiara Papetti, Marta Paterno, Massimo Ponti, Lorenzo Zane, and Leonardo Congiu. A population genomics insight by 2b-RAD reveals populations' uniqueness along the Italian coastline in *Leptopsammia pruvoti* (Scleractinia, Dendrophylliidae). *Diversity and Distributions*, 25(7):1101–1117, 2019. *Genetic sequence data archived under NCBI BioProject Accession*: PRJNA521302.

[15] Brendan H. Cornwell. Gene flow in the anemone *Anthopleura elegantissima* limits signatures of local adaptation across an extensive geographic range. *Molecular Ecology*,

29(14):2550–2566, 2020. *Genetic sequence data archived under NCBI BioProject Accession*: PRJNA632874.

[16] Nicole L Crane, Juliette Tariel, Jennifer E Caselle, Alan M Friedlander, D Ross Robertson, and Giacomo Bernardi. Clipperton Atoll as a model to study small marine populations: Endemism and the genomic consequences of small population size. *PLoS One*, 13(6):e0198901, 2018. *Genetic sequence data archived under NCBI BioProject Accession*: PRJNA448430, PRJNA448438.

[17] Hayley M. DeHart, Leocadio Blanco-Bercial, Mollie Passacantando, Jennifer M. Questel, and Ann Bucklin. Population connectivity of chaetognath *Eukrohnia hamata*. *National Center for Biotechnology Information*, BioProject Accession: PRJNA576675, Registration date: 09-October-2019.

[18] M. Lisette Delgado, Konrad Górski, Evelyn Habit, and Daniel E. Ruzzante. The effects of diadromy and its loss on genomic divergence: The case of amphidromous *Galaxias maculatus* populations. *Molecular Ecology*, 28(24):5217–5231, 2019. *Genetic sequence data archived under NCBI BioProject Accession*: PRJNA564121.

[19] James L. Dimond, Sanoosh K. Gamblewood, and Steven B. Roberts. Genetic and epigenetic insight into morphospecies in a reef coral. *Molecular Ecology*, 26(19):5031–5042, 2017. *Genetic sequence data archived under NCBI BioProject Accession*: PRJNA379028.

[20] R Fernández, M Schubert, AM Vargas-Velázquez, A Brownlow, GA Víkingsson, U Siebert, LF Jensen, N Øien, D Wall, E Rogan, et al. A genomewide catalogue of single nucleotide polymorphisms in white-beaked and Atlantic white-sided dolphins. *Molecular Ecology Resources*, 16(1):266–276, 2016. *Genetic sequence data archived under NCBI BioProject Accession*: PRJNA280898.

[21] María José Frugone, María Eugenia López, Nicolás I. Segovia, Theresa L. Cole, Andrew Lowther, Pierre Pistorius, Gisele P.M. Dantas, Maria Virginia Petry, Francesco Bonadonna, Phil Trathan, Andrea Polanowski, Barbara Wienecke, Ke Bi, Cynthia Y. Wang-Claypool, Jonathan M. Waters, Rauri C.K. Bowie, Elie Poulin, and Juliana A. Vianna. More than the eye can see: Genomic insights into the drivers of genetic differentiation in Royal/Macaroni penguins across the Southern Ocean. *Molecular Phylogenetics and Evolution*, 139:106563, 2019. *Genetic sequence data archived under NCBI BioProject Accession*: PRJNA523574.

[33] Sho Hosoya, Shotaro Hirase, Kiyoshi Kikuchi, Kusuto Nanjo, Yohei Nakamura, Hiroyoshi Kohno, and Mitsuhiko Sano. Random pcr-based genotyping by sequencing technology

GRAS-Di (genotyping by random amplicon sequencing, direct) reveals genetic structure of mangrove fishes. *Molecular Ecology Resources*, 19(5):1153–1163, 2019. *Genetic sequence data archived under NCBI BioProject Accession*: PRJDB7819.

[42] N. S. Locatelli and J. A. Drew. Population structure and clonal prevalence of scleractinian corals (*Montipora capitata* and *Porites compressa*) in Kaneohe Bay, Oahu. *bioRxiv*, 2019. *Genetic sequence data archived under NCBI BioProject Accession*: PRJNA544861.

[49] Marta Paterno, Marcello Schiavina, Giorgio Aglieri, Jamila Ben Souissi, Elisa Boscari, Renato Casagrandi, Aurore Chassanite, Mariachiara Chiantore, Leonardo Congiu, Giuseppe Guarnieri, Claudia Kruschel, Vesna Macic, Ilaria A. M. Marino, Chiara Papetti, Tomaso Patarnello, Lorenzo Zane, and Paco Melià. Population genomics meet Lagrangian simulations: Oceanographic patterns and long larval duration ensure connectivity among *Paracentrotus lividus* populations in the Adriatic and Ionian seas. *Ecology and Evolution*, 7(8):2463–2479, 2017. *Genetic sequence data archived under NCBI BioProject Accession*: PRJNA373825.

[50] Joseph B. Pfaller, Adam C. Payton, Karen A. Bjorndal, Alan B. Bolten, and Stuart F. McDaniel. Hitchhiking the high seas: Global genomics of rafting crabs. *Ecology and Evolution*, 9(3):957–974, 2019. *Genetic sequence data archived under NCBI BioProject Accession*: PRJNA471559.

[51] L. V. Plough. Population genomic analysis of the blue crab *Callinectes sapidus* using genotyping-by-sequencing. *Journal of Shellfish Research*, 36(1):249–261, 2017. *Genetic sequence data archived under NCBI BioProject Accession*: PRJNA340157.

[52] D. S. Portnoy, J. B. Puritz, C. M. Hollenbeck, J. Gelsleichter, D. Chapman, and J. R. Gold. Selection and sex-biased dispersal in a coastal shark: the influence of philopatry on adaptive variation. *Molecular Ecology*, 24(23):5877–5885, 2015. *Genetic sequence data archived under NCBI BioProject Accession*: PRJNA286089.

[53] Gregory N. Puncher, Alessia Cariani, Gregory E. Maes, Jeroen Van Houdt, Koen Herten, Rita Cannas, Naiara Rodriguez-Ezpeleta, Aitor Albaina, Andone Estonba, Molly Lutcavage, Alex Hanke, Jay Rooker, James S. Franks, Joseph M. Quattro, Gualtiero Basilone, Igaratza Fraile, Urtzi Laconcha, Nicolas Goñi, Ai Kimoto, David Macías, Francisco Alemany, Simeon Deguara, Salem W. Zgozi, Fulvio Garibaldi, Isik K. Oray, Firdes Saadet Karakulak, Noureddine Abid, Miguel N. Santos, Piero Addis, Haritz Arrizabalaga, and Fausto Tinti. Spatial dynamics and mixing of bluefin tuna in the Atlantic Ocean and Mediterranean Sea revealed using next-generation sequencing. *Molecular Ecology Resources*, 18(3):620–638, 2018. *Genetic sequence data archived under NCBI BioProject Accession*: PRJNA432036.

[54] Gregory Neils Puncher, Sherrylynn Rowe, George A. Rose, Nathalie M. Leblanc, Geneviève J. Parent, Yanjun Wang, and Scott A. Pavey. Chromosomal inversions in the Atlantic cod genome: Implications for management of Canada's Northern cod stock. *Fisheries Research*, 216:29–40, 2019. *Genetic sequence data archived under NCBI BioProject Accession*: PRJNA528403.

[55] Catherine M Purcell, Arun S Seetharam, Owyn Snodgrass, Sofia Ortega-García, John R Hyde, and Andrew J Severin. Insights into teleost sex determination from the *Seriola dorsalis* genome assembly. *BMC Genomics*, 19:1–11, 2018. *Genetic sequence data archived under NCBI BioProject Accession*: PRJNA319656.

[56] Jonathan B Puritz, John R Gold, and David S Portnoy. Fine-scale partitioning of genomic variation among recruits in an exploited fishery: Causes and consequences. *Scientific Reports*, 6(1):36095, 2016. *Genetic sequence data archived under NCBI BioProject Accession*: PRJNA329407.

[57] Hannah G. Reich, Sheila A. Kitchen, Kathryn H. Stankiewicz, Meghann Devlin-Durante, Nicole D. Fogarty, and Iliana B. Baums. Genotypic similarity among algal symbionts corresponds to associations with closely related coral hosts. *bioRxiv*, 2020. *Genetic sequence data archived under NCBI BioProject Accession*: PRJNA473816.

[58] Pablo Saenz-Agudelo, Joseph D. Dibattista, Marek J. Piatek, Michelle R. Gaither, Hugo B. Harrison, Gerrit B. Nanninga, and Michael L. Berumen. Seascape genetics along environ-

mental gradients in the Arabian Peninsula: Insights from ddRAD sequencing of anemonefishes. *Molecular Ecology*, 24(24):6241–6255, 2015. *Genetic sequence data archived under NCBI BioProject Accession*: PRJNA294760.

[75] Patricia H. Wepfer, Yuichi Nakajima, Makamas Sutthacheep, Veronica Z. Radice, Zoe Richards, Put Ang, Tullia Terraneo, Mareike Sudek, Atsushi Fujimura, Robert J. Toonen,

Alexander S. Mikheyev, Evan P. Economo, and Satoshi Mitarai. Evolutionary biogeography of the reef-building coral genus *Galaxea* across the Indo-Pacific Ocean. *Molecular Phylogenetics and Evolution*, 151:106905, 2020. *Genetic sequence data archived under NCBI BioProject Accession*: PRJNA576132.

[76] Shengyong Xu, Na Song, Linlin Zhao, Shanshan Cai, Zhiqiang Han, and Tianxiang Gao. Genomic evidence for local adaptation in the ovoviviparous marine fish *Sebastiscus marmoratus* with a background of population homogeneity. *Scientific Reports*, 7(1):1562, 2017. *Genetic sequence data archived under NCBI BioProject Accession*: PRJNA359404.

[77] Tyler S Zemlak, Evelyn M Habit, Sandra J Walde, Cecilia Carrea, and Daniel E Ruzzante. Surviving historical Patagonian landscapes and climate: molecular insights from *Galaxias maculatus*. *BMC Evolutionary Biology*, 10:1–18, 2010. *Genetic sequence data archived under NCBI BioProject Accession*: PRJNA564121.

[78] Bai-Dong Zhang, Dong-Xiu Xue, Yu-Long Li, and Jin-Xian Liu. RAD genotyping reveals fine-scale population structure and provides evidence for adaptive divergence in a commercially important fish from the Northwestern Pacific Ocean. *PeerJ*, 7:e7242, 2019. *Genetic sequence data archived under NCBI BioProject Accession*: PRJNA533707.

[79] Yunfeng Zhao, Wenzhu Peng, Huayang Guo, Baohua Chen, Zhixiong Zhou, Jian Xu, Dianchang Zhang, and Peng Xu. Population genomics reveals genetic divergence and adaptive differentiation of Chinese sea bass (*Lateolabrax maculatus*). *Marine Biotechnology*, 20:45–59, 2018. *Genetic sequence data archived under NCBI BioProject Accession*: PRJNA356786.

### Appendix S4: List of sequencing adapters used in sequence read cleaning

Below is a list of all adapter sequences that we searched for on the 3' end of all raw sequence reads downloaded from the International Nucleotide Sequence Database Collaboration. Lines that start with > denote adapter names and the directly succeeding line contains the adapter sequence.

We generated the below list from:
https://github.com/stephenturner/adapters/blob/master/adapters_combined_152_unique.fasta;
FastQC (https://github.com/s-andrews/FastQC/blob/master/Configuration/contaminant_list.txt);
and BBMap (https://github.com/BioInfoTools/BBMap/blob/master/resources/adapters.fa);
all accessed on April 9, 2020.

We removed duplicate adapters among the above sources and merged adapters with highly similar sequences using IUPAC base pair ambiguity (names of merged adapters begin with "MERGED"). Using the program cutadapt, we searched each FASTQ sequence file in our study for each of the below sequences. Datasets that contained adapters were easy to distinguish from those that did not; datasets with adapters returned hits in >25% of the reads per individual compared to <5% of reads per individual in clean datasets. When present, we removed identified adapter sequences from all reads in the dataset. We then performed a final adapter search to ensure no adapters remained in raw reads (i.e., all sequences below returned hits in $\sim$ 5% or fewer of the sequence reads per individual across the dataset).

---

```
>Adapter1 / aka PhiX_read1_adapter aka the Illumina common adapter
AGATCGGAAGAGCGGTTCAGCAGGAATGCCGAGACCGATCTCGTATGCCGTCTTCTGCTTG
>MERGED-Primers_and_dimers-GROUP
AATGATACGGCGACCACCGAGATCTACACTCTTTCCCTACACGACGCTCTTCCGATCTMRWKMDRMRGMSVSSWYMVVVRBBHWY
VCBVWBWCCSWWCWCGWMKSYCKTCYKMTSYTTG
>MERGED-PCR_Primer1_rc-PhiX_read2_adapter
AGATCGGAAGAGCGTCGTGTAGGGAAAGAGTGTAGATCTCGGTGGTCGCCGTATCATTAAAAAA
>MERGED-Reverse_adapter-1-Reverse_adapter-2
MKRTCKSWWRWRCACACGTCTGAACTCCAGTCACATCACGATCTCGTATGCCGTCTTCTGCTTG
>MERGED-TruSeq_Adapter_Index-GROUP1
GATCGGAAGAGCACACGTCTGAACTCCAGTCACVNBNVNNNATCTCGTATGCCGTCTTCTGCTTG
>MERGED-TruSeq_Adapter_Index-GROUP2
GATCGGAAGAGCACACGTCTGAACTCCAGTCACNNNNNNNNTCTCGTATGCCGTCTTCTGCTTG
>MERGED-TruSeq3_IndexedAdapter / PE2_rc-Bisulfite_R1
AGATCGGAAGAGCACACGTCTGAACTCCAGTCAC
>MERGED-ABI_Solid3_Adapter_B-ABI_Solid3_5p_AMP_Primer
CCACTACGCCTCCGCTTTCCTCTCTATGGGCAGTCGGTGAT
>MERGED-Illumina_DpnII_and_NlaIII_Adapters-GROUP
ACAGGTTCAGAGTTCTACAGTCCGACATG
>MERGED-I7_Primer_Nextera_XT-GROUP
CCGAGCCCACGAGACNNNNNNNNATCTCGTATGCCGTCTTCTGCTTG
>MERGED-Clontech_SMARTer_II_A_Oligonucleotide-Clontech_SMART_CDS_Primer_II_A
AAGCAGTGGTATCAACGCAGAGTACT
>MERGED-ABI_Solid3_Adapter_A-ABI_Solid3_3p_AMP_Primer
CTGCCCCGGGTTCCTCATTCTCTCAGCAGCATG
>MERGED-I5_Primer_Nextera_XT-GROUP
GACGCTGCCGACGANNNNNNNNGTGTAGATCTCGGTGGTCGCCGTATCATT
>MERGED-Bisulfite_R2-PE1_rc / TruSeq3_UniversalAdapter
AGATCGGAAGAGCGTCGTGTAGGGAAAGAGTGTA
>MERGED-Nextera_Transposase-GROUP
CTGTCTCTTATACACATCTSHSRBBSSVSDVGVVAGGC
>MERGED-Illumina_PCR_Primer_Index-GROUP
CAAGCAGAAGACGGCATACGAGATNNNNNNGTGACTGGAGTTC
```

```
>PrefixPE_1 / PE1
TACACTCTTTCCCTACACGACGCTCTTCCGATCT
>TruSeq2_PE_r
AGATCGGAAGAGCGGTTCAGCAGGAATGCCGAG
>I5_Adapter_Nextera
CTGATGGCGCGAGGGAGGCGTGTAGATCTCGGTGGTCGCCGTATCATT
>I7_Adapter_Nextera_No_Barcode
CTGAGCGGGCTGGCAAGGCAGACCGATCTCGTATGCCGTCTTCTGCTTG
>Nextera_LMP_Read1_External_Adapter / Illumina Multiplexing Index Sequencing Primer
GATCGGAAGAGCACACGTCTGAACTCCAGTCAC
>Nextera_LMP_Read2_External_Adapter
GATCGGAAGAGCGTCGTGTAGGGAAAGAGTGT
>Stop_Oligo_STP_8
CCACGGGAACGTGGTGGAATTC
>TruSeq2_SE
AGATCGGAAGAGCTCGTATGCCGTCTTCTGCTTG
>PrefixNX_1 / >PrefixNX/2
AGATGTGTATAAGAGACAG
>Illumina_Single_End_Adapter_1
GATCGGAAGAGCTCGTATGCCGTCTTCTGCTTG
>Illumina_Single_End_Adapter_2 / Illumina Single End PCR Primer 2 /
>Illumina DpnII expression Adapter 2
CAAGCAGAAGACGGCATACGAGCTCTTCCGATCT
>Illumina_Single_End_Sequencing_Primer / Illumina Paired End Adapter 1 /
>Illumina Paried End Sequencing Primer 1 / Illumina Multiplexing Adapter 2 /
>Illumina Multiplexing Read1 Sequencing Primer
ACACTCTTTCCCTACACGACGCTCTTCCGATCT
>Illumina_Paired_End_Adapter_2
GATCGGAAGAGCGGTTCAGCAGGAATGCCGAG
>Illumina_Paired_End_PCR_Primer_2 / PrefixPE/2 / PCR_Primer2
CAAGCAGAAGACGGCATACGAGATCGGTCTCGGCATTCCTGCTGAACCGCTCTTCCGATCT
>Illumina_Paired_End_Sequencing_Primer_2
CGGTCTCGGCATTCCTGCTGAACCGCTCTTCCGATCT
>Illumina_DpnII_expression_Adapter_2 / Illumina DpnII expression PCR Primer 1 /
>Illumina NlaIII expression Adapter 2 / Illumina NlaIII expression PCR Primer 1 /
>Illumina Small RNA RT Primer / Illumina DpnII Gex Adapter 2 /
>Illumina DpnII Gex PCR Primer 1 / Illumina NlaIII Gex Adapter 2.01 /
>Illumina NlaIII Gex PCR Primer 1 / Illumina Small RNA PCR Primer 1
CAAGCAGAAGACGGCATACGA
>Illumina_DpnII_expression_PCR_Primer_2 /
>Illumina NlaIII expression PCR Primer 2 / Illumina DpnII Gex PCR Primer 2 /
>Illumina NlaIII Gex PCR Primer 2
AATGATACGGCGACCACCGACAGGTTCAGAGTTCTACAGTCCGA
>Illumina_DpnII_expression_Sequencing_Primer / Illumina DpnII Gex Sequencing Primer
CGACAGGTTCAGAGTTCTACAGTCCGACGATC
>Illumina_NlaIII_expression_Sequencing_Primer / Illumina NlaIII Gex Sequencing Primer
CCGACAGGTTCAGAGTTCTACAGTCCGACATG
>Illumina_Multiplexing_Adapter_1
GATCGGAAGAGCACACGTCT
>Illumina_Multiplexing_PCR_Primer_2_01 / Illumina Multiplexing Read2 Sequencing Primer
GTGACTGGAGTTCAGACGTGTGCTCTTCCGATCT
>Illumina_DpnII_Gex_Adapter_1
GATCGTCGGACTGTAGAACTCTGAAC
>Illumina_DpnII_Gex_Adapter_2_01 / Illumina NlaIII Gex Adapter 2.02
TCGTATGCCGTCTTCTGCTTG
>Illumina_NlaIII_Gex_Adapter_1_01
TCGGACTGTAGAACTCTGAAC
>ABI_Dynabead_EcoP_Oligo
CTGATCTAGAGGTACCGGATCCCAGCAGT
>ABI_Solid3_EF1_alpha_Sense_Primer
CATGTGTGTTGAGAGCTTC
>ABI_Solid3_EF1_alpha_Antisense_Primer
GAAAACCAAAGTGGTCCAC
>ABI_Solid3_GAPDH_Forward_Primer
TTAGCACCCCTGGCCAAGG
>ABI_Solid3_GAPDH_Reverse_Primer
CTTACTCCTTGGAGGCCATG
>Clontech_Universal_Primer_Mix_Short
```

```
CTAATACGACTCACTATAGGGC
>Clontech_Universal_Primer_Mix_Long
CTAATACGACTCACTATAGGGCAAGCAGTGGTATCAACGCAGAGT
>FlowCell2
TTTTTTTTTTCAAGCAGAAGACGGCATACGA
>FlowCell1
TTTTTTTTTTAATGATACGGCGACCACCGAGATCTACAC
>Trans1
TCGTCGGCAGCGTCAGATGTGTATAAGAGACAG
>Trans2
GTCTCGTGGGCTCGGAGATGTGTATAAGAGACAG
```

### Appendix S5: Citations for biotic trait data

This appendix contains the citations for all of the biotic trait values used in this study. The citation numbers in this document correspond to the numbers in the columns labeled: "**<trait_name>_citation**" in the data file: "**/data/biotic/final_biotic_traits_for_suppmat-with-citation-numbers.csv**" in the Dryad archive for this study. Note that the trait Benthicity ("isBenthic") does not have a corresponding citation column (in "final_biotic_traits_for_suppmat-with-citation-numbers.csv") because these values reflect common knowledge within our author group. Other individual trait values that have "NA" in the corresponding citation column also reflect common knowledge within our author group.

We obtained as many trait values as possible from public trait databases (e.g., FishBase (R. Froese, D. Pauly, Editors 2023) and SeaLifeBase (M. L. D. Palomares, D. Pauly. Editors 2023)). For species/trait combinations not present in centralized databases (which were the majority), we performed Boolean searches in Web of Science or Google Scholar (e.g., <species name> AND egg size).

[6] BirdLife International. *Aythya marila*. *The IUCN Red List of Threatened Species 2018*, e.T22680398A132525108, 2018.

[7] E. O'Reilly, B.M. Titus, M.W. Nelsen, S. Ratchford, and N.E. Chadwick. Giant ephemeral anemones? Rapid growth and high mortality of corkscrew sea anemones *Bartholomea an-*

*nulata* (Le Sueur, 1817) under variable conditions. *Journal of Experimental Marine Biology and Ecology*, 509:44–53, 2018.

[8] D.A. Ebert. Reproductive biology of skates, *Bathyraja* (Ishiyama), along the eastern Bering Sea continental slope. *Journal of Fish Biology*, 66:618–649, 2005.

[9] M.R. Gaither, G.A. Gkafas, M. de Jong, F. Sarigol, F. Neat, T. Regnier, D. Moore, D.R. Gröcke, N. Hall, X. Liu, J. Kenny, A. Lucaci, M. Hughes, S. Haldenby, and A.R. Hoelzel. Genomics of habitat choice and adaptive evolution in a deep-sea fish. *Nature Ecology & Evolution*, 2:680–687, 2018.

[10] H.-J.T. Hoving, V.V. Laptikhovsky, and B.H. Robison. Vampire squid reproductive strategy is unique among coleoid cephalopods. *Current Biology*, 25:R322–R323, 2015.

[11] E.M. Salas, G. Bernardi, M.L. Berumen, M.R. Gaither, and L.A. Rocha. RADseq analyses reveal concordant Indian Ocean biogeographic and phylogeographic boundaries in the reef fish *Dascyllus trimaculatus*. *Royal Society Open Science*, 6:172413, 2019.

[12] B.A. Fach. Modeling the influence of hydrodynamic processes on anchovy distribution and connectivity in the Black Sea. *Turkish Journal of Fisheries and Aquatic Sciences*, 14, 2014.

[13] BirdLife International. *Eudyptes chrysolophus*. *The IUCN Red List of Threatened Species 2020*, e.T22697793A184720991, 2020.

[14] D. Grawunder, E.A. Hambleton, M. Bucher, I. Wolfowicz, N. Bechtoldt, and A. Guse. Induction of gametogenesis in the cnidarian endosymbiosis model *Aiptasia sp. Scientific Reports*, 5, 2015.

[15] A.J. Edwards. FAO species catalogue. Vol. 10. Gadiform fishes of the world (Order Gadiformes). An annotated and illustrated catalogue of cods, hakes, grenadiers and other gadiform fishes known to date. *Marine Pollution Bulletin*, 24:326–327, 1990.

[16] D. Bowen. *Halichoerus grypus*. *The IUCN Red List of Threatened Species 2016*, e.T9660A45226042, 2016.

[25] G. Braulik, G. Minton, M. Amano, and A. Bjørge. *Phocoena phocoena*. *The IUCN Red List of Threatened Species 2020*, e.T17027A50369903, 2020.

[26] L. Rojas-Bracho and B.L. Taylor. *Phocoena sinus*. *The IUCN Red List of Threatened Species 2017*, e.T17028A50370296, 2017.

[27] P. Gèlin, C. Fauvelot, V. Mehn, S. Bureau, H. Rouzè, and H. Magalon. Superclone expansion, long-distance clonal dispersal and local genetic structuring in the coral *Pocillopora damicornis* type $\beta$ in Reunion Island, south western Indian Ocean. *PLOS ONE*, 12:e0169692, 2017.

[28] K. Soong. Sexual reproductive patterns of shallow-water reef corals in Panama. *Bulletin of Marine Science*, 49:832–846, 1991.

[29] BirdLife International. *Pygoscelis adeliae*. *The IUCN Red List of Threatened Species 2020*, e.T22697758A157660553, 2020.

[30] BirdLife International. *Pygoscelis antarcticus*. *The IUCN Red List of Threatened Species 2020*, e.T22697761A184807209, 2020.

[31] BirdLife International. *Pygoscelis papua*. *The IUCN Red List of Threatened Species 2020*, e.T22697755A157664581, 2020.

[32] N. Pasiec. *Rhizophora mangle* (red mangrove). In *CABI Compendium*. CABI Digital Library, 2015.

[33] J.P. Wourms. Reproduction and development of *Sebastes* in the context of the evolution of piscine viviparity. In *Rockfishes of the genus Sebastes: Their reproduction and early life history*, pages 111–126. Springer, 1991.

[34] D. Moran, C.K. Smith, B. Gara, and C.W. Poortenaar. Reproductive behaviour and early development in yellowtail kingfish (*Seriola lalandi* Valenciennes 1833). *Aquaculture*, 262:95–104, 2007.

[35] A.L. Gould, K.E. Dougan, S.T. Koenigbauer, and P.V. Dunlap. Life history of the symbiotically luminous cardinalfish *Siphamia tubifer* (Perciformes: Apogonidae). *Journal of Fish Biology*, 89:1359–1377, 2016.

[36] C. Bartilotti and A. Dos Santos. The secret life of deep-sea shrimps: Ecological and evolutionary clues from the larval description of *Systellaspis debilis* (Caridea: Oplophoridae). *PeerJ*, 7:e7334, 2019.

[37] BirdLife International. *Uria aalge*. *The IUCN Red List of Threatened Species 2018*, e.T22694841A132577296, 2018.

[38] F. Trillmich. *Zalophus wollebaeki*. *The IUCN Red List of Threatened Species 2015*, e.T41668A45230540, 2015.

[39] M.E. Bushnell, J.T. Claisse, and C.W. Laidley. Lunar and seasonal patterns in fecundity of an indeterminate, multiple-spawning surgeonfish, the yellow tang *Zebrasoma flavescens*. *Journal of Fish Biology*, 76:1343–1361, 2010.

[40] J.A.B. Claydon. *The structure and dynamics of spawning aggregations of coral reef fish*. PhD thesis, James Cook University, 2005.

[41] G.V. Clucas, J.L. Younger, D. Kao, L. Emmerson, C. Southwell, B. Wienecke, A.D. Rogers, C. Bost, G.D. Miller, M.J. Polito, et al. Comparative population genomics reveals key barriers to dispersal in Southern Ocean penguins. *Molecular Ecology*, 27:4680–4697, 2018.

[42] G.R. Hoff. Identification of multiple nursery habitats of skates in the eastern Bering Sea. *Journal of Fish Biology*, 88:1746–1757, 2016.

[43] J.H. Churchill, J.P. Kritzer, M.J. Dean, J.H. Grabowski, and G.D. Sherwood. Patterns of larval-stage connectivity of Atlantic cod (*Gadus morhua*) within the Gulf of Maine in relation to current structure and a proposed fisheries closure. *ICES Journal of Marine Science*, 74:20–30, 2017.

[44] J.M. González-Irusta and P.J. Wright. Spawning grounds of Atlantic cod (*Gadus morhua*) in the North Sea. *ICES Journal of Marine Science: Journal du Conseil*, 73:304–315, 2015.

[45] A. Ellison, E. Farnsworth, and G. Moore. *Laguncularia racemosa*. *The IUCN Red List of Threatened Species 2010*, e.T178798A7609219, 2010.

[46] B.A. McIntyre, E.E. McPhee-Shaw, M.B.A. Hatch, and S.M. Arellano. Location matters: Passive and active factors affect the vertical distribution of Olympia oyster (*Ostrea lurida*) larvae. *Estuaries and Coasts*, 44:199–213, 2020.

[47] L.E. Timm, T.L. Jackson, J.A. Browder, and H.D. Bracken-Grissom. Population genomics of the commercially important Gulf of Mexico pink shrimp *Farfantepenaeus duorarum* (Burkenroad, 1939) support models of juvenile transport around the Florida peninsula. *Frontiers in Ecology and Evolution*, 9, 2021.

[60] J.S. Nelson, T.C. Grande, and M.V.H. Wilson. *Fishes of the world*. John Wiley & Sons, Hoboken, New Jersey, 2016.

[72] R.T. Hanlon and J.B. Messenger. *Cephalopod Behaviour*. Cambridge University Press, 1998.

[73] S.W. Davies, M.E. Strader, J.T. Kool, C.D. Kenkel, and M.V. Matz. Modeled differences of coral life-history traits influence the refugium potential of a remote Caribbean reef. *Coral Reefs*, 36:913–925, 2017.

[74] J.S. Barber, J.E. Dexter, S.K. Grossman, C.M. Greiner, and J.T. Mcardle. Low temperature brooding of Olympia oysters (*Ostrea lurida*) in northern Puget Sound. *Journal of Shellfish Research*, 35:351–357, 2016.

[75] C. Carreras, A. Garcìa-Cisneros, O.S. Wangensteen, V. Ordóñez, C. Palacín, M. Pascual, and X. Turon. East is east and west is west: Population genomics and hierarchical analyses reveal genetic structure and adaptation footprints in the keystone species *Paracentrotus lividus* (Echinoidea). *Diversity and Distributions*, 26:382–398, 2019.

[76] J.M. Leis. The pelagic stage of reef fishes: The larval biology of coral reef fishes. In *The Ecology of Fishes on Coral Reefs*, pages 183–230. Springer, 1991.

[77] M.F. Strathmann. *Reproduction and Development of Marine Invertebrates of the Northern Pacific Coast*. University of Washington Press, 1987.

[87] T.D. Williams. *The Penguins Spheniscidae. Bird Familes of the World*, volume 2. Oxford University Press, 1995.

[88] I. Wolfowicz, S. Baumgarten, P.A. Voss, E.A. Hambleton, C.R. Voolstra, M. Hatta, and A. Guse. *Aiptasia sp.* larvae as a model to reveal mechanisms of symbiont selection in cnidarians. *Scientific Reports*, 6, 2016.

[89] D. Macedo, I. Caballero, M. Mateos, R. Leblois, S. McCay, and L.A. Hurtado. Population genetics and historical demographic inferences of the blue crab *Callinectes sapidus* in the US based on microsatellites. *PeerJ*, 7:e7780, 2019.

[90] COSEWIC. *COSEWIC assessment and status report on the Roundnose Grenadier Coryphaenoides rupestris in Canada*. Committee on the Status of Endangered Wildlife in Canada, 2008.

[105] M. Bucher, I. Wolfowicz, P.A. Voss, E.A. Hambleton, and A. Guse. Development and symbiosis establishment in the cnidarian endosymbiosis model *Aiptasia sp*. *Scientific Reports*, 6, 2016.

[106] R.C. Babcock and A.J. Heyward. Larval development of certain gamete-spawning scleractinian corals. *Coral Reefs*, 5:111–116, 1986.

[107] B.D. Limer, J. Bloomberg, and D.M. Holstein. The influence of eddies on coral larval retention in the Flower Garden banks. *Frontiers in Marine Science*, 7, 2020.

[124] J.B. Puritz, J.R. Gold, and D.S. Portnoy. Fine-scale partitioning of genomic variation among recruits in an exploited fishery: Causes and consequences. *Scientific Reports*, 6, 2016.

[134] C.D. Storlazzi, M. van Ormondt, Y.-L. Chen, and E.P.L. Elias. Modeling fine-scale coral larval dispersal and interisland connectivity to help designate mutually-supporting coral reef marine protected areas: Insights from Maui Nui, Hawaii. *Frontiers in Marine Science*, 4, 2017.

[135] L.E. Timm, H.D. Bracken-Grissom, A. Sosnowski, M. Breitbart, M. Vecchione, and H. Judkins. Population genomics of three deep-sea cephalopod species reveals connectivity between the Gulf of Mexico and northwestern Atlantic ocean. *Deep Sea Research Part I: Oceanographic Research Papers*, 158:103222, 2020.

[136] B. Villamor, M. Bernal, and C. Hernandez. Models describing mackerel (*Scomber scombrus*) early life growth in the north and northwest of the Iberian Peninsula in 2000. *Scientia Marina*, 68:571–583, 2004.

[137] M.S. Love, M. Yoklavich, and L.K. Thorsteinson. *The Rockfishes of the Northeast Pacific*. University of California Press, 2002.

[138] A.L. Gould and P.V. Dunlap. Shedding light on specificity: Population genomic structure of a symbiosis between a coral reef fish and luminous bacterium. *Frontiers in Microbiology*, 10, 2019.

[152] E. Beukhof, T.S. Dencker, M.L.D. Palomares, and A. Maureaud. A trait collection of marine fish species from North Atlantic and Northeast Pacific continental shelf seas [dataset]. *Pangaea*, 2019.

[159] J.E.N. Veron and M. Stafford-Smith. *Corals of the World*. Australian Institute of Marine Science, 2000.

[172] C.L. Hunter. *Genotypic Diversity and Population Structure of the Hawaiian Reef Coral, Porites compressa*. PhD thesis, University of Hawai'i, Honolulu, 1988.

[179] A.V. Golikov, F.R. Ceia, R.M. Sabirov, J.D. Ablett, I.G. Gleadall, G. Gudmundsson, H.J. Hoving, H. Judkins, J. Pálsson, A.L. Reid, R. Rosas-Luis, E.K. Shea, R. Schwarz, and Xavier J.C. The first global deep-sea stable isotope assessment reveals the unique trophic ecology of vampire squid *Vampyroteuthis infernalis* (Cephalopoda). *Scientific Reports*, 9, 2019.

[180] T. Foster and J. Gilmour. Egg size and fecundity of biannually spawning corals at Scott Reef. *Scientific Reports*, 10, 2020.

[191] K. De Baets, C. Klug, D. Korn, and N.H. Landman. Data from: Early evolutionary trends in ammonoid embryonic development, 2015. *Dryad*, 10.5061/dryad.k84c3k8j.

[199] M.E. Bushnell. *Reproduction of Zebrasoma flavescens: oocyte maturation, spawning patterns, and an estimate of reproductive potential for female yellow tang in Hawaii*. PhD thesis, University of Hawaii at Manoa, 2007.

[200] D.A. Pavlov and N.G. Emel'yanova. Fertility of ovulated oocytes after their short-term storage in water in several species of marine tropical fishes. *Journal of Ichthyology*, 48:655–664, 2008.

[201] B.B. Collette. Hemiramphidae, halfbeaks. In K.E. Carpenter and N. De Angelis, editors, *The living marine resources of the Eastern Central Atlantic. Volume 3: Bony fishes part 1 (Elopiformes to Scorpaeniformes)*. FAO Species Identification Guide for Fishery Purposes, Rome, 2016.

[212] P. Casale and A.D. Tucker. *Caretta caretta* (amended version of 2015 assessment). *The IUCN Red List of Threatened Species 2017*, e.T3897A119333622, 2017.

[213] Stazione zoologica di Napoli. *Fauna e Flora del Golfo di Napoli, Monografia 38: Uova, Larve e Stadi Giovanili di Teleostei*. Springer-Verlag, 1892. Monographs elaborated with the use of material collected and serialized by S. Lo Bianco from U. D'Ancona, L. Sanzo, A. Sparta, F. Bertolini, G. Montalenti, E. Padoa, S. Ranzi, E. Tortonese, M. Vialli.

### Appendix S6: Search parameters and citations for Global Biodiversity Information Facility species occurrence records

This appendix contains the search parameters and citations for the resulting occurrence records that we downloaded from the Global Biodiversity Information Facility (GBIF) for each species included in this study. The search parameters section denotes the exact search and filtering terms that we passed to the `occ_download` function (rgbif R package V3.5.0, Chamberlain and Boettiger 2017) to retrieve species occurrence records from GBIF for each species in this study. The citations capture the specific snapshots of GBIF data used in this study; the DOIs are specific to the exact search parameters that we used and the date that we executed the query.

##### Search and filtering parameters used to retrieve GBIF records:

```
library(rgbif)

OurSearchRequest <- occ_download(type='and',
                                 pred('taxonKey',taxonKey),
                                 pred("hasGeospatialIssue", FALSE),
                                 pred("hasCoordinate", TRUE),
                                 pred_not(
                                   pred("basisOfRecord", "FOSSIL_SPECIMEN")
                                 ),
                                 pred_or(
                                   pred_not(pred("establishmentMeans","MANAGED")),
                                   pred_not(pred_notnull("establishmentMeans"))
                                 ),
                                 pred_or(
                                   pred("occurrenceStatus","PRESENT"),
                                   pred_not(pred_notnull("occurrenceStatus"))
                                 ),
                                 format='SIMPLE_CSV',
                                 user=user,pwd=password,email=email)
```
